# Molecular architecture and spatial organisation of proteasomes in the human sperm nucleus

**DOI:** 10.64898/2025.12.16.694293

**Authors:** Piotr Kolata, Ália dos Santos, Oliver Knowles, Tom Dendooven, Matteo Allegretti

## Abstract

Proteasomes are proteolytic machines essential for proteostasis and genome integrity. They consist of a 20S core particle capped by regulatory complexes that confer substrate selectivity (Voges, 1999). Despite evidence that proteasome function is necessary for successful spermatogenesis and fertilisation (Sutovsky, 2011; Xiong *et al*., 2022), the presence, composition, and role of proteasomes within the sperm nucleus have remained a subject of debate. Here, we used *in situ* electron cryo-tomography (cryo-ET) in human sperm cells to elucidate the molecular architecture of nuclear proteasomes, which cluster in DNA-free, nuclear cavities within the sperm nucleus. Subtomogram averaging revealed that the main population of proteasomes consists of 20S core particles, with a smaller fraction of 20S capped by PA200 activator, solved at sub-nanometre resolution. Subsequent single-particle cryo-EM analysis of purified native human sperm proteasomes further elucidated atomic features of the essential testis-specific subunit α4s, showing the presence of a unique splice variant. Notably, our reconstruction resolved a native peptide in the catalytic β2 subunit, providing novel insights into the proteolysis mechanism and PA200-mediated enhancement of trypsin activity. Finally, we show nuclear enrichment of proteasomes during sperm cell differentiation in human testis tissue, with 20S and PA200 clustering following meiosis, at the spermatid stage. Our findings shed light on the unique organisation and compositional diversity of nuclear proteasomes in human sperm cells, disclosing novel molecular insights into their catalytic function.

## Introduction

The proteasome is an essential multi-catalytic protease complex consisting of a barrel-shaped 20S core particle (CP) of ∼750 kDa, capped by regulatory particles. In eukaryotes, the 20S core comprises 28 subunits arranged in four stacked rings (α1–7, β1–7)_2_, forming a hollow cylinder (da Fonseca *et al*., 2012). The outermost rings of α-subunits create two gated pores that control substrate entry, while the inner β-rings harbour the protease active sites. To degrade proteins, the 20S CP associates with proteasome activators that recognise substrates and open the pore to enable their entry. The main activator, the 19S regulatory particle, associates with the 20S to form a 26S holoenzyme. Apart from opening the gate, the 19S activator unfolds ubiquitylated proteins in an ATP-dependent manner and threads them into the 20S CP for proteolysis (Dong et al., 2019).

The proteasome plays critical roles in both cytoplasmic and nuclear protein quality control (Enenkel *et al*., 2022; Guo, 2022). In the nucleus, proteasomes are crucial for degrading cell-cycle regulators, clearing misfolded proteins, and dismantling protein complexes at DNA repair foci (Guo, 2022; Lafarga M, 2002). The nuclear proteasome pool is dynamically regulated. Under acute cellular stress, such as heat shock, oxidative or osmotic stress, nutrient starvation, or inhibition of proteasome activity or nuclear export, proteasomes can reorganise into intranuclear, membraneless condensates via ubiquitin-mediated liquid–liquid phase separation (Enenkel *et al*., 2022; Lee *et al*., 2021; Uriarte *et al*., 2021; Yasuda *et al*., 2020).

In the context of reproduction, a growing body of evidence indicates that spermatozoa harbour enzymatically active proteasomes that are essential both during spermatogenesis and after fertilisation (Xiong *et al*., 2022). Consistent with this, inhibition of proteasome function leads to a dose-dependent reduction in motility (Hackerova *et al*., 2023) and disrupts sperm-cell capacitation (Kong *et al*., 2009; Morales *et al*., 2003; Morales *et al*., 2004; Zigo *et al*., 2019). Proteasomes have been detected in multiple compartments of the sperm cell, including the acrosome, periacrosomal surface, plasma membrane, perinuclear theca (Morales *et al*., 2004; Rawe *et al*., 2008; Song *et al*., 2021; Sutovsky, 2011; Sutovsky *et al*., 2004; Zhang *et al*., 2022) as well as the connecting piece and tail (Rawe *et al*., 2008; Zimmerman and Sutovsky, 2009). Studies across different mammalian model organisms propose a role for sperm proteasomes in the degradation of glycoproteins in the egg’s zona pellucida (ZP), and show that inhibition of proteasome activity blocks sperm penetration, thereby preventing fertilisation (Saldivar-Hernandez *et al*., 2015; Sutovsky, 2011; Sutovsky *et al*., 2004; Zimmerman and Sutovsky, 2009).

While the cytosolic, extracellular and membrane-associated functions of sperm proteasomes have been well studied (Sutovsky, 2011; Xiong *et al*., 2022), less is known about the role of nuclear proteasomes in mammalian sperm. Proteasomes are essential for nuclear DNA compaction during spermatogenesis, where the histones in the chromatin of post-meiotic germ cells are replaced with protamines. The timely removal of histones is facilitated by a testis-specific proteasome variant, in which the canonical α4 (PMSA7) proteasome subunit is replaced by the α4s (PSMA8) subunit in spermatocytes (Zhang *et al*., 2019; Zhang *et al*., 2021; Zivkovic *et al*., 2022). α4s appears to be essential for effective meiotic progression, being localised to, and potentially involved in the disassembly of the synaptonemal complex (Gomez *et al*., 2019; Zhang *et al*., 2021; Zivkovic *et al*., 2022). In spermatid nuclei, the PA200 activator engages the testis-specific proteasome, allowing acetylation-dependent proteolysis of core histones (Jiang *et al*., 2021; Qian *et al*., 2013; Sato *et al*., 2023; Ustrell, 2002). Moreover, both PA200 and α4s are thought to be essential for sperm differentiation and maturation. In mice lacking PA200, the removal of core histones from elongating spermatids is significantly delayed, while deletion of the α4s subunit leads not only to defective histone removal, but also _to meiotic arrest and male infertility_ (Gomez *et al*., 2019; Khor *et al*., 2006; Xiong *et al*., 2022; ^Zhang *et al*., 2019; Zhang *et al*., 2021)^. How these nuclear proteasomes are organised in the dense sperm head remains unknown, but proteasome components were found to co-localise to electron-bright regions of the sperm nucleus (“lacunae”), using immunogold labelling and transmission electron microscopy (TEM) (Haraguchi *et al*., 2007). The size of these nuclear *lacunae* has been proposed to correlate with levels of chromatin condensation, sperm motility and fertility outcomes (Chemes and Alvarez Sedo, 2012; Tanaka et al., 2012).

Here, we used in situ cryo-electron tomography (cryo-ET) and sub-tomogram averaging to reveal large proteasome populations within nuclear *lacunae* of mature and motile human sperm cells. The *lacunae* were up to 1 µm in length and were enriched in 20S proteasomes, with an additional fraction of PA200-capped proteasomes. Using immunohistochemistry in human testis tissue, we show that proteasome-rich *lacunae* develop in post-meiotic cells at the spermatid stage. In addition, our high-resolution structures of native human sperm proteasomes revealed a unique splice variant of the testis-specific subunit α4s and an unexpected density in the β2 subunit active site. Collectively, our work maps and characterises proteasome populations in the sperm nucleus, shedding light on substrate engagement and PA200-driven proteasome activation mechanisms.

## Results

### Nuclear lacunae in human sperm cells contain three populations of proteasome complexes

To investigate the localisation and ultrastructure of nuclear proteasomes in human sperm cells, we used cryo-ET. Human sperm cells were subjected to cryo-focused ion beam (cryo-FIB) milling to generate thin lamellae (Fig. 1a). In the lamellae maps, we found electron-bright regions in the sperm nuclei, similar to the *lacunae* described by Haraguchi et al (Haraguchi et al., 2007). Reconstructed tomograms of these regions (Fig. 1a) revealed that *lacunae* were segregated from the condensed protamine-DNA, as membrane-less compartments, and enriched in barrel-shaped protein complexes (Fig. 1b, Extended Data Fig. 1a). Sub-tomogram averaging and 3D classification of these complexes revealed three different proteasome populations: 20S, 20S-PA200 and PA200-20S-PA200. We obtained a 6.8 Å map of the 20S proteasome CP (94.4% of assigned particles), a 9.5Å map of the 20S capped by a single PA200 activator (20S-PA200, 5.3% particles), and a 14.4 Å map of the 20S capped by two PA200 activators (PA200-20S-PA200, 0.3% particles) (Fig. 1c-d, Extended Data Fig. 1b-c, Extended Data Video 1). Next, we investigated whether nuclear proteasome *lacunae* were a conserved feature between species. For this, we performed cryo-ET on lamellae of mouse sperm cells (Extended Data Fig. 2a-b). Similar to human sperm cells, we found proteasome complexes in the nuclear *lacunae* of mouse sperm cells (Extended Data Fig. 2b). However, the size of the proteasomal *lacunae* was significantly smaller compared to human (Extended Data Fig. 1, Extended Data Fig. 2a-b).

**Fig. 1.**
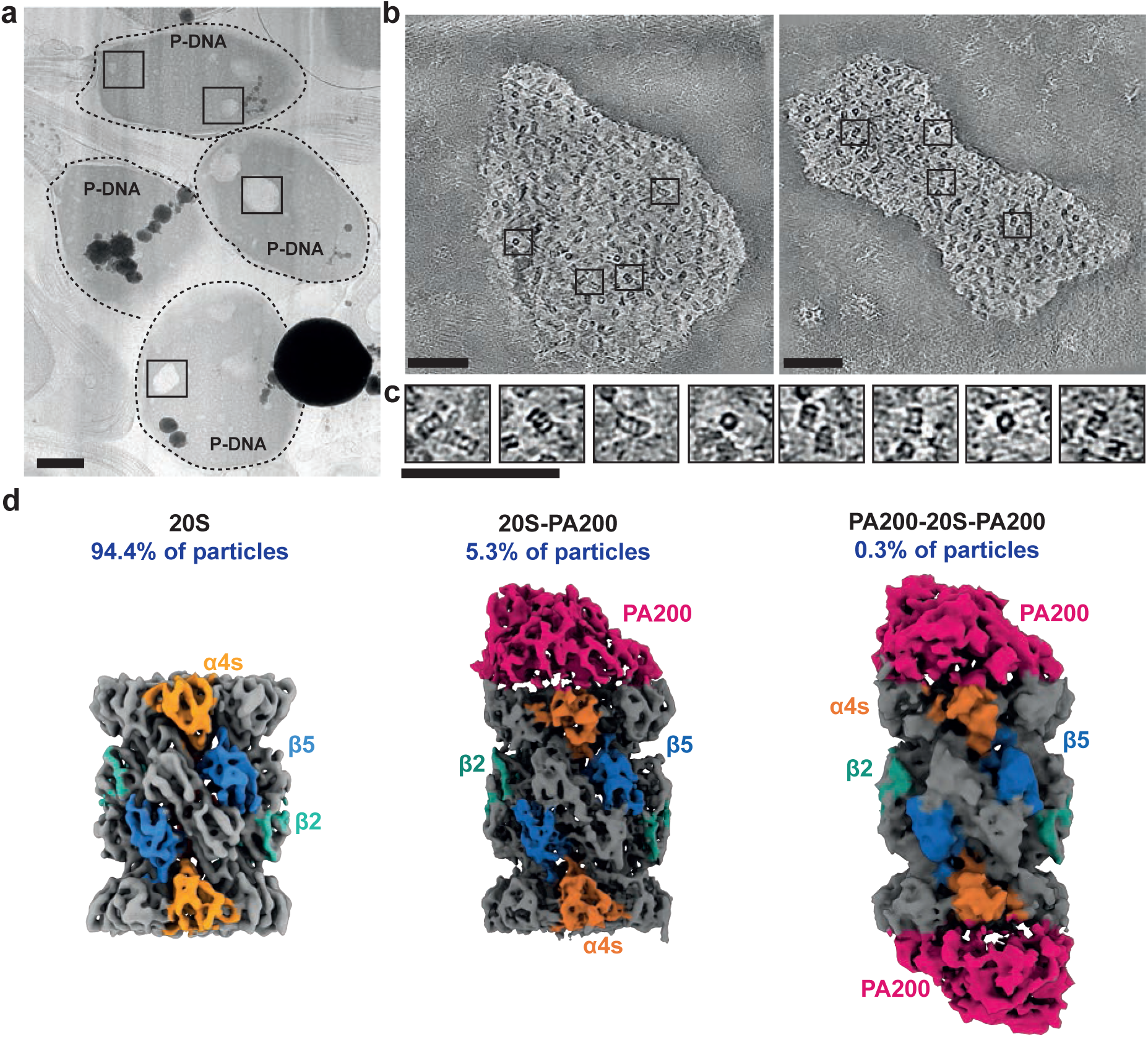
Nuclear lacunae in human sperm cells are proteasomal compartments. **a**, cryo-TEM low magnification lamellae map showing four human sperm heads. Arrows indicate regions with sperm lacunae targeted for tilt-series acquisition. Scale bar = 1 *μ*m. **b,** z-slices from reconstructed tomograms showing barrel-shaped protein complexes inside membrane-less nuclear compartments. Scale bar = 100nm. **c,** zoom-in from tomograms shown in **b**, showing barrel-shaped particles inside *lacunae*. Scale bar = 100 nm. **d,** subtomogram averaging classes showing three populations: 20S complexes (43,128 particles, 94.4% of total particles); 20S-PA200 complexes (2,430 particles, 5.3% of total particles); and PA200-20S-PA200 (245 particles, 0.3% of total particles).

Lamellae prepared for cryo-ET (∼ thickness 120-220 nm) only represent 5% of the total volume of a sperm cell. To assess the size and variability of proteasome-rich *lacunae*, we performed volumetric imaging of entire human sperm cells using cryo-FIB/SEM (Fig. 2a, Extended Data Video 2) (Schertel *et al*., 2013). The reconstructed volumes reveal that nuclear *lacunae* (Fig. 2a-b) have an average Feret diameter - defined as the longest linear distance of a segmented *lacuna* - of 326 ± 6 nm (Fig. 2c-d). Among all segmented human sperm *lacunae,* 7% exceeded a Feret diameter of 500 nm, and 2.3% were larger than 1 *μ*m, consistent with our cryo-ET observations (Fig. 1a, Extended Data Fig. 1a, Fig. 2c). Similarly, segmented cryo-FIB/SEM volumes of mouse sperm cells showed a marked reduction in the average Feret diameter of *lacunae* to 173 nm ± 2 nm, with only 0.2% larger than 500 nm, none exceeding 600 nm and 42% smaller than 150 nm. (Fig. 2c-d, Extended Data Fig. 2b-c).

**Fig. 2.**
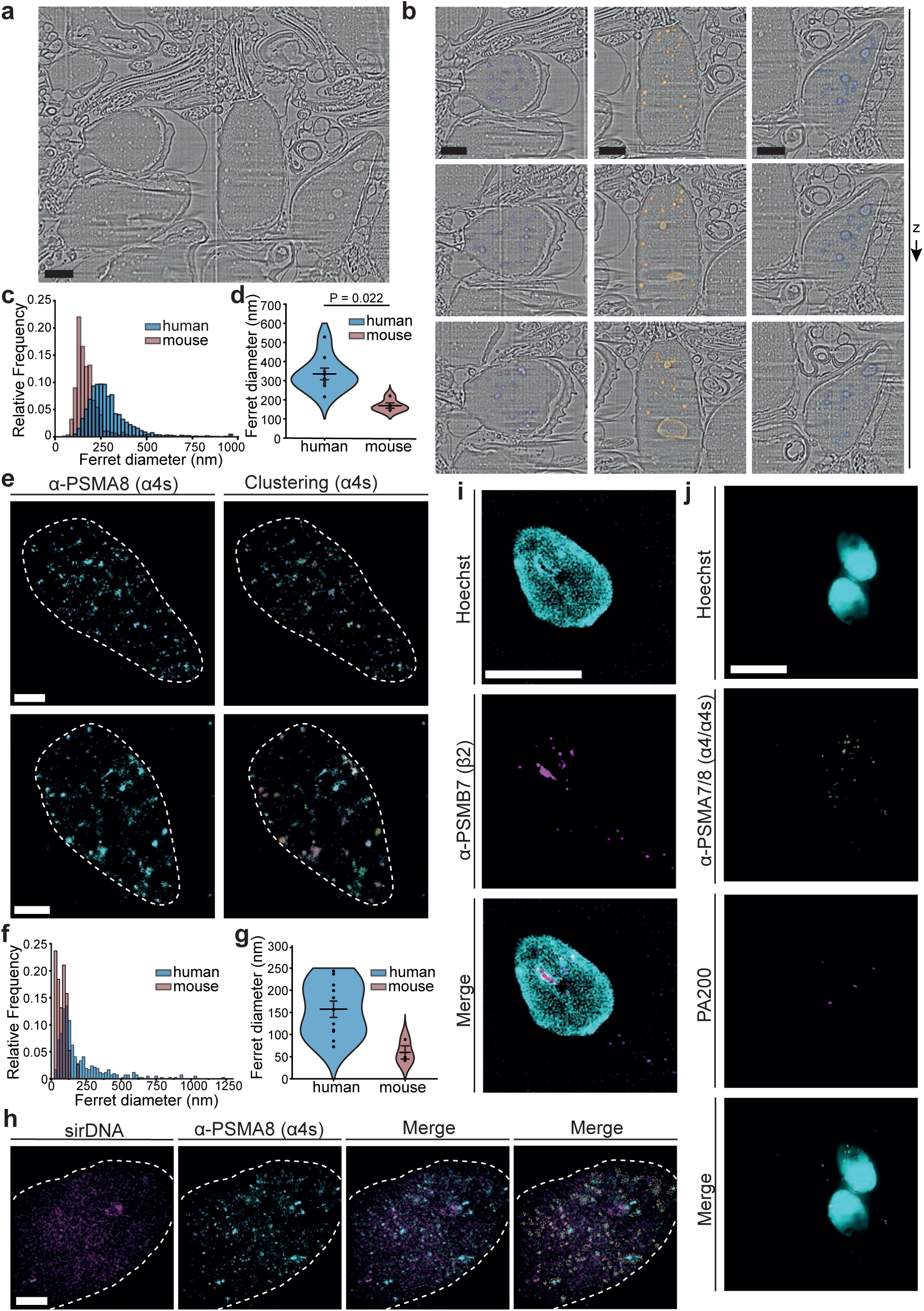
Human sperm nuclear *lacunae* proteasomes are positive for testis-specific subunit PSMA8, segregate from DNA and are variable in size. **a**, z-slice of 3D reconstructed cryo-scanning electron microscopy volumetric data of human sperm cells. Sperm lacunae are visible in the nucleus. Scale = 1 *μ*m. **b,** detail of three human sperm heads from volumetric data shown in **a**. Segmented lacunae are shown in colour (purple, yellow and blue) at three different z heights. **c,** Relative frequency plot of Feret diameter of segmented human sperm lacunae (blue) and comparison with values from mouse data (red, Extended Data Fig. 2). Feret diameter was defined as the longest xy distance of a segmented lacuna. The histogram depicts the values for 2,147 segmented lacunae segmented from 9 human sperm cells, and 866 lacunae from 5 mouse sperm cells. Human *lacunae* mean Feret diameter ± standard error of the mean (s.e.m.) = 326 ± 6 nm. Mouse *lacunae* mean Feret diameter ± s.e.m = 173.3 ± 2.1 nm. **d,** Violin plots for data shown in **d**. Each data point represents the average lacunae diameter of an individual cell. Mean ± s.e.m. diameter is plotted for human (blue, 335.3 ± 30.5 nm, n = 9 cells) and mouse cells (red, 169.4 ± 13.6 nm, n = 5 cells). Statistical analysis: t-test, *P* = 0.0022. **e,** Stochastic Optical Reconstruction Microscopy (STORM) of sperm-specific proteasome subunit ***α***4s (PSMA8) in human sperm cells. White dashed line shows sperm nuclear region. Panels on the right show ***α***4s cluster identification following DBSCAN analysis of ***α***4s within nuclear region. Each cluster is shown in a different colour. Scale bars = 1*μ*m. **f,** Histogram of ***α***4s clusters identified using STORM in human (blue) and mouse cells (red, Extended Data Fig. 3a). Histogram represents values from 415 ***α***4s clusters obtained from 11 human sperm cells and 37 ***α***4s clusters from 3 mouse sperm cells. Human proteasome clusters mean Feret diameter ± standard error of the mean (s.e.m.) = 199.0 ± 7.2 nm. Mouse proteasome clusters mean Feret diameter ± s.e.m = 72.0 ± 5.9 nm. **g,** Violin plots of data shown in **f**. Data points represent average Feret diameter of ***α***4s clusters in each cell. Mean ± s.e.m. diameter is shown for human (blue, 157.3 ± 18.4 nm, n = 11 cells) and mouse cells (red, 59.7 ± 25.7 nm, n = 3 cells). For mouse data > 10 cells were measured by STORM, but clusters were only identified in 3 cells. Clusters were identified in all measured human cells. **h,** STORM imaging of human sperm cell DNA using SiR-DNA and immunofluorescence staining against ***α***4s. Regions of ***α***4s density segregated from DNA are shown within dashed yellow lines. Scale bar = 1*μ*m. **i,** Structured Illumination Microscopy (SIM) showing proteasome subunit PSMB7 (*β*2) forming foci in the nucleus of sperm cells (magenta) in regions of no DNA staining (Hoechst). Scale bar = 5 *μ*m. **j,** SIM imaging of human sperm cells showing nuclear foci of *α*4/*α*4s (PSMA7/8) proteasome subunit (green) and partial co-localisation with PA200 (magenta). Small *α*4/*α*4s foci are also present in the tail. Scale bar = 5 *μ*m.

To corroborate these results with an orthogonal method and measure the size distribution of nuclear proteasome clusters (rather than *lacunae*), we used Stochastic Optical Reconstruction Microscopy (STORM), followed by cluster analysis with Density-Based Spatial Clustering of Applications with Noise (DBSCAN) (Ram *et al*., 2010). The sperm-specific 20S subunit PSMA8 (*α*4s) formed clusters within the nucleus of human sperm cells (Fig. 2e), displaying cluster size distributions comparable to those obtained with cryo-SEM volumetric data, although with smaller predicted dimensions (Fig. 2f-g). Human sperm *α*4s clusters had an average Feret diameter of 199 nm, with 7.2% exceeding 500 nm, and 0.7% larger than 1 *μ*m. Furthermore, proteasome clusters did not colocalise with DNA staining, consistent with our cryo-ET observation that *lacunae* are segregated from DNA (Fig. 2h). Mouse sperm STORM data were also in line with cryo-SEM segmentation data, showing fewer clusters than in human sperm and a marked reduction in cluster size, with an average Feret diameter of 72 nm, 97% smaller than 150 nm and none exceeding a diameter of 200 nm (Fig. 2f-g; Extended Data Fig. 3a). In mouse sperm cells, PSMA8 (*α*4s) localisations did not always form regions of sufficient density to constitute clusters, measured by DBSCAN. Immunofluorescence staining of human sperm cells, followed by Structured Illumination Microscopy (SIM), confirmed the presence of proteasome foci localised to regions largely devoid of DNA, as shown by both STORM (Fig. 2h) and SIM (Fig. 2i-j, Extended Data Fig. 3d). A small subset of 20S foci co-localised with PA200 (Fig. 2j), consistent with the low fraction of capped proteasomes identified with subtomogram averaging (Fig. 1d). SIM imaging of mouse sperm cells revealed a similar distribution of 20S and PA200 (Extended Data Fig. 3b). Interestingly, we did not detect any signal for 19S (PSMC1) in the nucleus of either human or mouse sperm cells. Instead, 19S was enriched in the tail region, especially in human samples (Extended Data Fig. 3c-d).

Together, our integrative imaging data, spanning cryo-ET, cryo-SEM and STORM imaging, demonstrate three populations of proteasome complexes clustering within nuclear *lacunae* of human spermatozoa: uncapped 20S CP, 20S-PA200 and PA200-20S-PA200, with no detectable quantities of 26S complexes. Human nuclear *lacunae* were also larger on average than those of mouse sperm, emphasising species-specific differences in gamete nuclear organisation.

### Single-particle cryo-EM reveals unique features of proteasomes isolated from human sperm cells

To investigate the structure of sperm-specific proteasomes, we purified native proteasomes from human sperm (Extended Data Fig. 4a). Sample purity was monitored with negative-stain EM and immunoblotting (Extended Data Fig. 4b-d), and cryo-EM micrographs (Extended Data Fig. 4e) allowed us to obtain a 1.85 Å map of the sperm-specific 20S CP (Fig. 3a-b, Extended Data Fig. 4e-f, Extended Data Fig. 5a, Extended Table 1, Extended Data Fig. 6a-f). The native sperm 20S proteasome did not contain immuno- or thymo-proteasome subunits. Specifically, we identified clear densities for the sidechains corresponding to the canonical β5 and β2 subunits, instead of the immunoproteasome-specific β5i and β2i subunits (Fig. 3c and Extended Data Fig. 6c-f). Notably, our structure not only confirms that α4 is replaced by α4s in sperm (Fig. 3c, Extended Data Fig. 6a, b) but also shows that the human sperm α4s subunit is an alternative splice variant (UniProt ID: Q8TAA3-5) that lacks the α4s-specific 77-82^α4s^ loop, referred to as α4s* hereafter (Fig. 3c). α4s* shares a near identical fold with the canonical α4 subunit (Extended Data Fig. 7a) (RMSD between Cα atoms of α4s* and α4 (PDB ID: 5LE5) (Schrader J., 2016) of 0.567Å). Accordingly, the overall fold of the sperm proteasome is highly conserved relative to the somatic form (RMSD between Cα atoms of an asymmetric unit of 0.538 Å). While the overall sequence conservation between the α4s* and α4 is very high (83.9 % identity), 68% of the substituted amino acids are solvent-exposed (Fig. 3d, Extended Data Fig. 7a). Most of these substitutions are conserved across mammals and render the surface of α4s* less negatively charged than that of α4 (Fig. 3d, dashed line). Inter-subunit contacts between α4s* or α4 remain unchanged, except for a conserved Ile82^α4^ to Val84^α4s*^ substitution at the α4s/α3 interface (Extended Data Fig. 6a-b; Extended Data Fig. 7b). Residues that interact with the proteasome assembly chaperone PAC4 were also preserved in α4s*, indicating that sperm 20S proteasome can also be assembled using the canonical machinery (Adolf *et al*., 2024; Zhang *et al*., 2024). Similar to the canonical 20S CP, the gate of the sperm 20S is fully ordered and adopts a constricted conformation. However, a conserved N-terminal Arg^α4s*^ insertion creates a basic patch at the inner pore surface (Extended Data Fig. 6a, Extended Data Fig. 7c), which may affect the kinetics of substrate entry or product release from the sperm proteasome. Finally, the active sites of β1, β2, and β5 subunits closely match those of the mature somatic 20S CP, with pro-peptides cleaved and catalytic residues in the same conformation (Extended Data Fig. 7d-f).

**Fig. 3.**
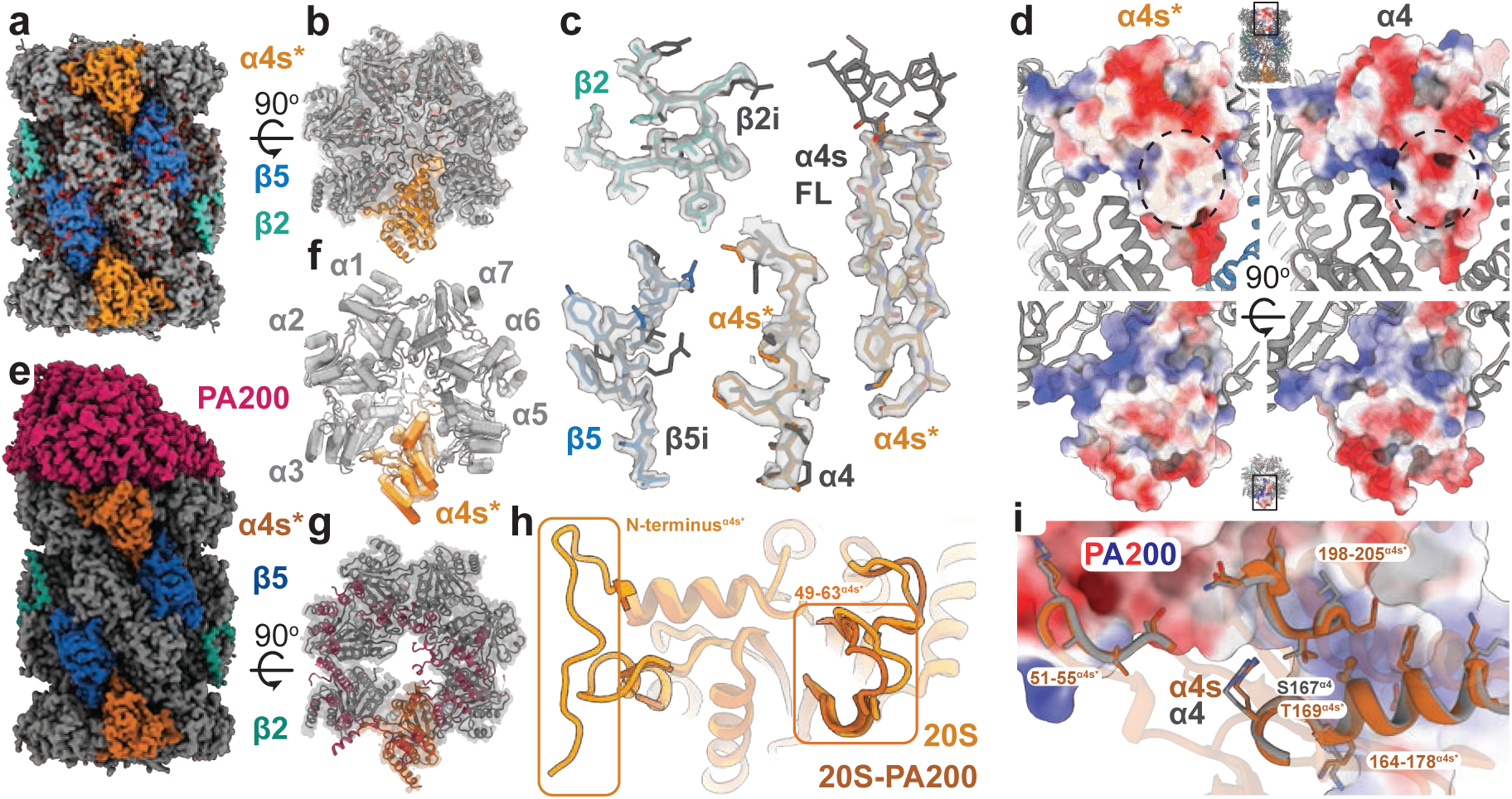
Single-particle cryo-EM structures of proteasomes isolated from human sperm cells. **a,** segmented map of a sperm 20S CP at 1.85 Å resolution. Key subunits are indicated: α4s* (orange), β5 (blue), β2 (cyan). **b**, constricted pore of the sperm 20S CP. The segmented map is shown as a transparent surface, and the corresponding structure is shown in a cartoon representation. The map and structure are coloured according to **a**. **c**, the canonical β2 and β5 subunits are present in the sperm proteasome, and an alternatively spliced α4s variant, referred to as α4s*, replaces the α4 subunit. Residues 119-127^β2/β2i^, 141-149 ^β5/β5i^, 217-224^α4s*^/215-222^α4^, as well as 63-77^α4s*^/63-83^α4s^ are shown as sticks and coloured as in **a**. The densities corresponding to these residue stretches in the sperm 20S core particle are shown as a transparent surface. β2, β5, and α4s coordinates from sperm 20S CP are shown. β2i and β5i immunoproteasome coordinates (PDB ID: 7b12) (Klein *et al*., 2021) were rigid-body fitted into the sperm 20S map for comparison. The canonical α4s (UniProt ID: Q8TAA3-1) is referred to as α4s FL. The alternatively spliced variant lacking the α4s-specific loop 77-82^α4s^ ^FL^ (UniProt ID: Q8TAA3-5) is referred to as α4s*. α4s FL coordinates were modelled as described in Materials and Methods. **d,** most substitutions in α4s occur at the solvent-accessible surface. α4s* (left) and α4 (right, PDB ID: 5LE5) coordinates are shown as an electrostatic surface, while other proteasomal subunits are shown as cartoons. The calculated mean coulombic electrostatic potential is of −0.09 kcal/mol·*e* for α4 and 0.14 kcal/mol·*e* for α4s^*.^ The substitutions Asp39^α4^ → Asn41^α4s^, Thr141^α4^ → Ile143^α4s^, Glu182^α4^ → Ala184^α4s^, Asp185^α4^ → Ser187^α4s^, and Lys227^α4^ → Leu229^α4s^ create an extensive, uncharged patch on the α4s* surface, shown as a dashed circle. Both side view (top) and top view (bottom) are shown. The shown regions of the structure are indicated in the insets by black rectangles. **e,** the segmented map of sperm 20S-PA200 proteasome at 2.9 Å resolution. Key subunits are indicated: PA200 (dark pink), α4s* (dark orange), β5 (dark blue), β2 (dark cyan). **f,** conformational changes in the α-subunits occurring upon PA200 engagement. The structures are shown as cartoons with α-helices as tubes. The 20S-PA200 structure is opaque in darker tones, while the 20S is transparent in brighter tones. α4s* is coloured in orange tones. **g,** PA200 binding opens the proteasome pore. The segmented map is shown as a transparent surface, and the corresponding structure is shown as cartoons. For PA200, only structural features interacting with the α-subunits are shown. **h,** conformational changes in α4s* upon binding of PA200. α4s* from the sperm 20S core particle is shown in orange, while the one from 20S-PA200 is in dark orange. The disordered terminus of 20S-PA200 and the 49-63^α4s*^ loop are marked with rectangles. **i,** Upon docking, α4s* interacts with PA200 via mostly conserved interface formed by loops 51-55^α4s*^, and 198-205^α4s*^, as well as with the 164-178^α4s*^ helix. α4 (grey) and α4s* (dark orange) structural features interacting with PA200 are indicated and shown as cartoons and sticks. PA200 of sperm 20S-PA200 is shown as an electrostatic surface. The mutation S167^α4^ → T169^α4s^ is indicated, with the T169^α4s^ side chain shown in ball-and-stick representation.

We also solved the structure of the native 20S-PA200 complex at 2.9 Å resolution (Fig. 3e-g; Extended Data Fig. 4a-f, Extended Data Fig. 5b, Extended Data Table 1). Overall, the structure closely resembled that of the recombinant somatic 20S-PA200 (PDB ID: 6KWY) (Guan *et al*., 2020) (RMSD between Cα atoms = 0.639 Å), with an open α-ring gate that allows substrate entry (Fig. 3b, f, g, h, Extended Data Fig. 8a-d). All α-subunits form extensive contacts with PA200-(Extended Data Fig. 8a-d). α4s* interacts with PA200 (Fig. 3i) via numerous, mostly polar contacts which are conserved between α4 and α4s*, except for a conserved Ser167^α4^ to Thr169^α4s*^ substitution (Fig. 3i, Extended Data Fig. 6a, Extended Data Fig. 8a). Given the similarity of the two residues, and the plethora of other interactions at the 20S/PA200 interface, we conclude that α4s* does not have an increased affinity for the PA200 activator.

We identified a small fraction of 26S proteasomes in the 20S-PA200 sample and solved their structure at ∼3.3 Å resolution (Extended Data Fig. 8e-g, Extended Data Fig. 5c, Extended Table 1). Since the 20S proteasome complex was not detected within nuclear *lacunae* (Fig. 1d), they likely originate from the sperm tail. The resolution of the 19S cap was lower (Extended Data Fig. 5c), due to its flexibility, but it was sufficient to fit the somatic 26S proteasome structure (PDB ID: 5L4G) (Schweitzer *et al*., 2016) into the density. The fit allowed us to preclude major structural rearrangements in the sperm 26S relative to the somatic variant. Correspondingly, the α4/α4s does not interact with the 19S cap in our map (Extended Data Fig. 8e-f).

### A tripeptide in the β2 subunit active site reveals insights into substrate recognition and PA200-mediated enhancement of trypsin catalytic activity

In the β2 subunit of the sperm 20S CP, we observed an unexpected, continuous, and branched density protruding from the nucleophilic Thr1^β2^ β-hydroxyl group towards the S1, S2, and S3 sites (Fig. 4a-c). We modelled it as an Arg-containing tripeptide (Fig. 4a-c). Since no additional polypeptide density extended downstream of the carbonyl carbon, the unknown density is either a trapped acyl-enzyme reaction intermediate following peptide cleavage, or a peptide-like inhibitor forming a hemiacetal (Huber *et al*., 2016). Regardless of its identity, the density reveals the coordination of a native-like substrate in the S1 pocket for the first time, to our knowledge. The Arg side chain is stabilised by the Asp53^β2^ carboxyl group, Ser32^β2^ carbonyl oxygen, as well as by a network of water molecules (Fig. 4c). S1 pocket broadening is required to engage the substrate. This is achieved through rearrangement of the Cys31^β2^ side chain (Fig. 4b, inset). In our map, the density allows modelling of the Cys31^β2^ side chain in both conformations, indicating that the substrate is present with partial occupancy (Fig. 4b). The remaining residues of the tripeptide bind the S2 and S3 pockets in a configuration similar to that seen in inhibitor-bound and propeptide-containing structures (Groll M., 1997; Huber *et al*., 2016; Schrader J., 2016) (Fig. 4a). Interestingly, the catalytic water that was proposed to perform a nucleophilic attack on the ester linkage is not positioned at an appropriate geometry in relation to the carbonyl group to resolve the acyl-enzyme intermediate (Fig. 4a) (Fleming, 2010). This likely explains trapping of the intermediate in the active site.

**Fig. 4.**
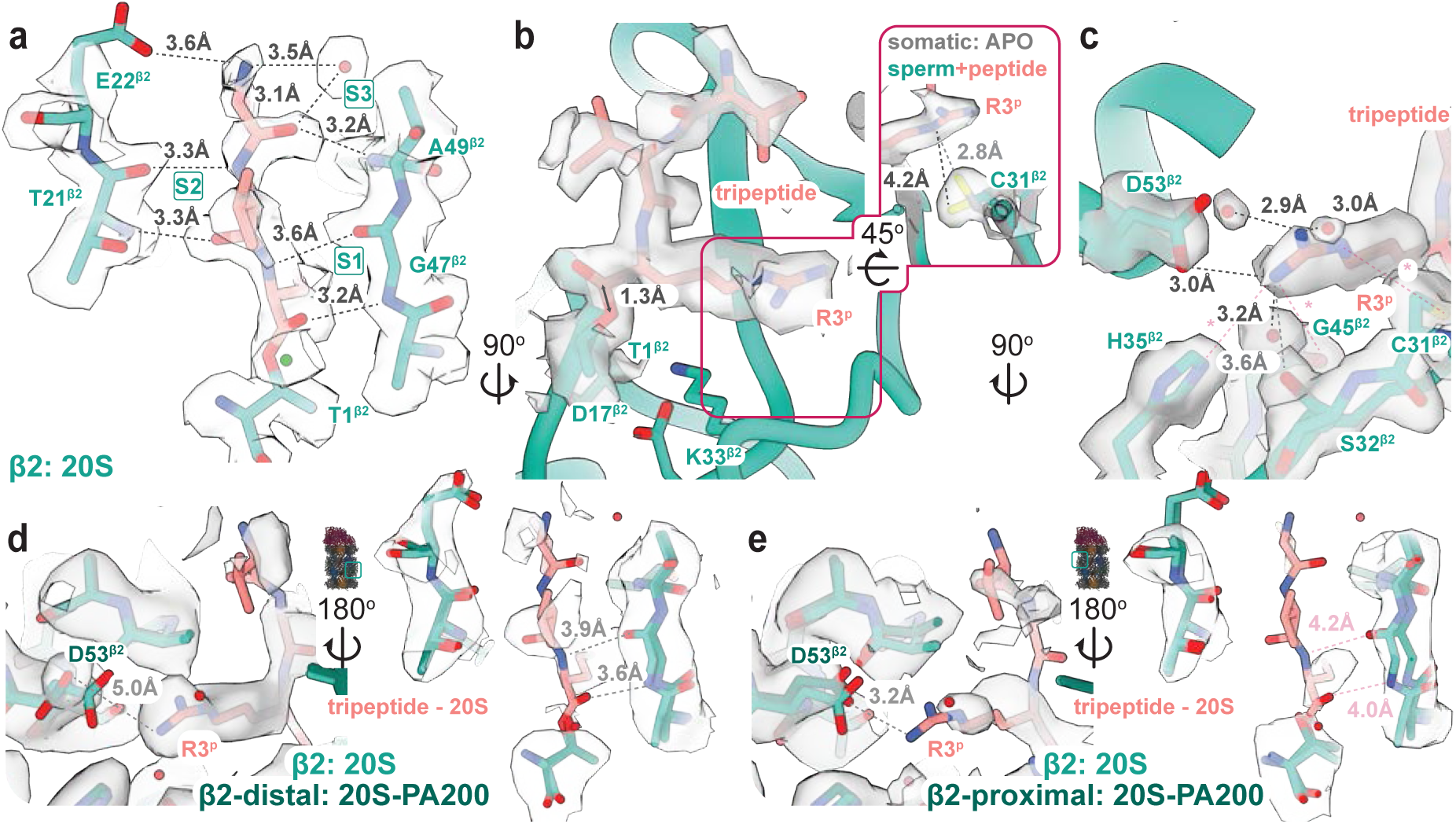
A peptide-like density in the β2 active site provides insights into substrate recognition and PA200-mediated enhancement of trypsin activity. **a,** the substrate tripeptide completes an antiparallel β-sheet in β2 subunit of sperm 20S core particle by forming hydrogen bonds with Thr21^β2^, Gly47^β2^, and Ala49^β2^. The residues coordinating the substrate polypeptide backbone (cyan), as well as the substrate itself (pink), are shown as sticks with the corresponding densities shown as transparent surfaces. The S1, S2, and S3 positions are indicated. The dashed lines indicate hydrogen bonding (distances shown). The water molecule, previously identified as a catalytic nucleophile for deacylation, is shown in green. **b,** density protruding from the Thr1^β2^ β-hydroxyl group is identified as a tripeptide acyl-enzyme intermediate. It contains a clear Arg density (R3^P^) with its putative carbonyl carbon ∼1.3 Å away from the nucleophilic oxygen, consistent with a single C-O covalent linkage (Allen, 2006). The tripeptide is shown as pink sticks, covalently linked to the catalytic Thr1^β2^ β-hydroxyl. The corresponding density is shown as a transparent surface. The catalytic residues Thr1^β2^, Asp17^β2^, and Lys33^β2^ are indicated and shown as cyan sticks. The β2 subunit is shown as cyan cartoons. The inset shows the Cys31^β2^ rearrangement required to incorporate the arginine side chain. In grey, the somatic 20S proteasome structure in the APO state is shown. The corresponding region in the sperm 20S CP is shown in cyan (the enzyme) and pink (the tripeptide). Cys31^β2^ and the tripeptide are shown as sticks, with their corresponding densities shown as transparent surfaces. The distance between the thiol group and N^δ^ of Arg is shown for both the sperm 20S CP (black dashed line) and the putative interaction in the APO somatic enzyme that would lead to a steric clash (grey dashed line). **c,** the coordination of substrate arginine side chain in the β2 active site of sperm 20S CP. The arginine, as well as its coordinating Asp53^β2^, Ser32^β2^, and water molecules, are shown as sticks with corresponding densities shown as transparent surfaces. Black/grey dashed lines indicate hydrogen bonds and salt bridges with distances shown. The distances between N^ω^ of arginine and His35^β2^ (4.0 Å), N^ω^ of arginine and Gly45^β2^ carbonyl (5.8 Å), N^δ^ of arginine and Cys31^β2^ (4.2 Å) preclude interactions and are indicated as red stars and dashed lines. **d,** the structure of β2 subunit distal from the PA200 activator (β2-distal) is conserved with the sperm 20S CP β2. The main difference is the apparent rearrangement of the D53^β2^ side chain away from the Arg side chain. The β2 subunits from the sperm 20S structure (cyan), β2-distal from 20S-PA200 structure (dark cyan), as well as the tripeptide from the 20S structure (pink), are shown as sticks. The corresponding 20S-PA200 density is shown as a transparent surface. The inset shows the position of the β2-distal in the 20S-PA200 structure. In the left panel, the same view as in **b** is used. The distance between the N^ω^ of arginine and the Asp53^β2^ side chain in the β2-distal is marked with a dashed line and indicated. The right panel shows conformational conservation of the substrate-coordinating S1, S2, and S3 sites between the sperm 20S β2 subunits and the β2-distal of the sperm 20S-PA200. The distances between the substrate backbone and S1 residues are marked with grey dashed lines and indicated. **e,** engaging PA200 causes conformational changes in the β2 subunit proximal to the activator (β2-proximal), expanding the substrate binding site to facilitate product release. The same features as in **d** are shown for the β2-proximal subunit of sperm 20S-PA200. The tripeptide from the 20S structure is shown for reference, as only residual substrate density is present in the β2-proximal site. The distances between the hypothetical substrate and the key coordinating residues are marked with dashed lines and indicated. In **d** and **e**, the map is shown at a 0.291 density threshold level. In all panels, amino acids are marked using single-letter amino acid code.

Engaging the PA200 cap increases the proteasome trypsin activity at least threefold (Toste Rego and da Fonseca, 2019; Ustrell, 2002). Aligning the 20S β2 subunit to the corresponding regions in the 20S-PA200 structure reveals the possible reasons for this (Fig. 4d-e). Particularly, the structure is almost entirely conserved with 20S in the activator-distal β2 subunit, including the presence of the peptide density (Fig. 4d). In turn, in the activator-proximal β2 subunit, the Asp53^β2^–containing helix is tilted away from the putative product, while the β-strands of S2 and S3 sites widen apart. These conformational changes likely reduce the coordination strength and allow efficient release of the product. In line with this, only the residual signal is detected in the S1 pocket of the activator-proximal β2, while the S2 and S3 positions are unoccupied (Fig. 4e), indicating an increased peptide processing rate. Altogether, our 20S structure reveals unprecedented insights into the binding of a native-like peptide into the trypsin catalytic site of the human proteasome and uncovers a distinct substrate recognition mechanism by β2. Moreover, the structure of the 20S-PA200 proteasome from a native source explains the molecular determinants of trypsin activity enhancement upon cap engagement.

### Human proteasome lacunae with 20S and PA200 form during spermatid stage of spermatogenesis

To investigate how the distribution and levels of proteasome complexes change during human sperm cell differentiation (Fig. 5a), we obtained healthy testis tissue from human donors and performed immunohistochemistry followed by confocal microscopy (Fig. 5b and Extended Data Fig. 9a-l). 20S subunits PSMA7/8 (α4/ α4s), PSMB7 (β2) and PSMA3 (α7) were present in all cells at the different stages of differentiation from spermatogonia to spermatozoa (Fig. 5b-c, Extended Data Fig. 9a-d). Interestingly, there was an enrichment in nuclear 20S core particles as differentiation progressed, with significantly higher levels in the nucleus of spermatocytes and spermatids, relative to spermatogonia (Fig. 3c and Extended Data Fig. 9b). Furthermore, large clusters of 20S were clearly present in the nuclei of both spermatids and mature spermatozoa in human tissue (Fig. 5c, stars in SPTD and SZ panels), which coincided with the presence of protamine in the nucleus (PRM1) (Extended Data Fig. 9k-l). These clusters, similar to what we described in ejaculated sperm (Fig. 2e-j), appeared to preferentially form in nuclear regions without DNA, i.e. *lacunae* (Figure 5c) (SD and SZ stars). In mouse testes, we observed a similar pattern, with significant enrichment in nuclear 20S CP as differentiation progressed from the spermatogonia to the spermatid stage (Extended Data Fig. 10a-c). However, proteasomal clustering within the nuclei of post-meiotic cells was markedly reduced compared to human tissue, particularly in mature spermatozoa (Extended Data Fig. 10b, SD and SZ panels, dashed lines). In somatic cells, PA200 is enriched in the nucleus, where it was shown to be involved in histone degradation (Jiang *et al*., 2021; Qian *et al*., 2013; Sato *et al*., 2023; Ustrell, 2002). Our human tissue data confirms the presence of PA200 in nuclei of both spermatogonia and differentiating sperm cells. In human spermatids and spermatozoa, PA200 formed clusters that co-localise with 20S (Extended Data Fig. 9a, Fig. 2j, Fig. 5c).

**Fig. 5.**
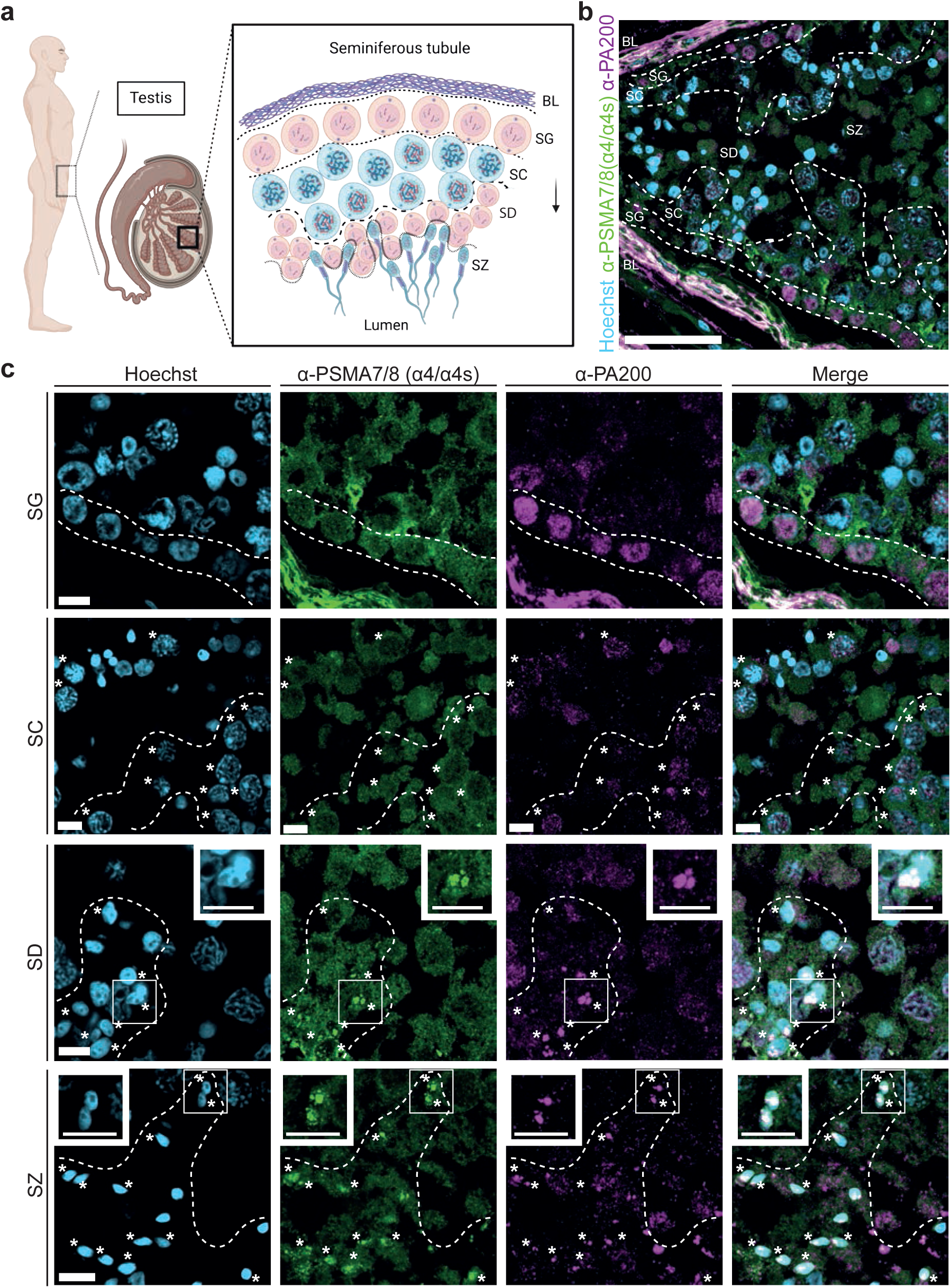
Proteasome reorganisation and clustering during human sperm cell differentiation. **a,** diagram showing the organisation of seminiferous tubules in human testes. Cells within seminiferous tubules are surrounded by a basal lamina (BL). Spermatogenesis progresses from the outer region of the tubule towards the central lumen. Spermatogonia (SG), localised close to the BL, undergo two rounds of meiosis. Spermatocytes (SC) – meiotic cells - occupy the median region of the tubules, and post-meiotic haploid spermatids (SD) lie close to the lumen, where they differentiate into mature spermatozoa (SZ). **b,** confocal microscopy image showing an overview of a seminiferous tubule from a healthy human donor. Immunohistochemistry staining shows distribution of proteasome subunit PSMA7/8 (*α*4/*α*4s) in green, proteasome adaptor PA200 in magenta and DNA (Hoechst in cyan). Individual channels for this image are shown in Extended Data Fig. 9a. White dashed lines represent boundaries between different cell types. Scale bar = 50 *μ*m **c,** Distribution of PSMA7/8 (*α*4/*α*4s) and PA200 at different differentiation stages in human testes. Panels represent zoomed-in regions of the overview shown in **b.** Each cell type (SG, SC, SD and SZ) is represented within white dashed line boundaries and/or by white stars. Top right insets in the SD and top left insets in the SZ panels represent zoomed-in regions of the areas inside the white rectangle in the respective panel. Scale bar = 5 *μ*m.

Next, we investigated the localisation of 19S in human tissue, as neither our light microscopy nor our cryo-ET data from ejaculated sperm cells showed the presence of this adaptor in the nucleus (Fig. 1d, Extended Data Fig. 4d). Our data show that 19S is highly enriched in the lumen of seminiferous tubules (Extended Data Fig. 9e-j) where it localises to acetylated tubulin in the tails of mature spermatozoa and the growing tail of spermatids (Extended Data Fig. 9f-j). Interestingly, we found that in Sertoli cells within human testis tissue, the 19S adaptor is enriched only in the nucleus (Extended Data Fig. 9e, f, h). We also observed localisation of 19S in spermatocyte cell protrusions that co-localised with acetylated tubulin and could represent primary cilia (Pérez-Moreno *et al*., 2025) (Extended Data Fig. 9g, orange stars). In mice, on the other hand, we detected a lower signal for 19S in the tail of mature spermatozoa (Fig. 4c) that we could not detect in differentiating spermatids/spermatozoa co-localising with acetylated tubulin (Extended Data Fig. 4c, Extended Data Fig. 10d).

Overall, our human tissue data reveal that clustering of both the 20S sperm proteasome and the activator PA200 in nuclear *lacunae* occurs from the spermatid stage, concurrent with DNA protamination. Conversely, in mouse tissue, clustering is minimal in spermatids and was not detected in spermatozoa using immunohistochemistry, further emphasising differences between the two species. Finally, 19S was mostly present in the tail of human spermatids and spermatozoa.

## Discussion

Previous studies indicate that nuclear proteasomal activity is indispensable for chromatin remodelling and sperm development (Qian *et al*., 2013). However, the presence, localisation and function of nuclear proteasomes in sperm cells have remained unexplored. The pioneering work of Haraguchi et al. (Haraguchi *et al*., 2007) first proposed the presence of proteasomes within nuclear cavities of both rat and human sperm heads. Chemes et al. further annotated these membrane-less cavities in sperm DNA as *lacunae* and linked them to dysregulated proteolytic activity (Chemes and Alvarez Sedo, 2012). Here, we show that human sperm nuclear *lacunae* are large assemblies of proteasome complexes, and we characterise their molecular architecture.

Subtomogram averaging of proteasomes *in situ* has previously revealed important aspects of cytoplasmic and nuclear proteasome biology (Albert *et al*., 2017; Albert *et al*., 2020; Guo *et al*., 2018). In this study, we used cryo-ET and sub-tomogram averaging to establish that nuclear sperm *lacunae* are populated by a mixture of 20S proteasomes and PA200-capped proteasomes (Fig. 1, Extended Data Fig. 1, Extended Data Fig. 2, Extended Data Video 1). Volume imaging and super-resolution STORM microscopy further revealed that proteasome-rich *lacunae* vary significantly in size and morphology (Fig. 2a-d; Extended Data Fig. 3b-c), and coincide with areas devoid of DNA, in agreement with our cryo-ET data (Fig. 2e-j). Furthermore, we show that nuclear proteasome clustering is conserved in mice but reveal species-specific differences by showing that mouse sperm *lacunae* are markedly smaller and fewer in number.

In the sperm proteasome, the canonical α4 subunit is replaced with the testis-specific α4s. This substitution is conserved across species and is essential for sperm differentiation and fertility (Zhang *et al*., 2019). Our near-atomic resolution cryo-EM structure of the 20S proteasome CP (Fig. 3a-d; Extended Data Fig. 5f; Extended Data Fig. 6; Extended Data Fig. 7) reveals the exclusive presence of a specific splice variant in human, which we term α4s*. The main amino-acid substitutions in α4s*, relative to α4, are located on the external surface of the 20S (Fig. 3d, Extended Data Fig. 6a-b; Extended Data Fig. 7a-b), suggesting a possible role in recruiting specific factors during spermatogenic differentiation. This would be consistent with reports that α4s deletion leads to meiotic arrest and data showing preferential interactomes of α4s proteasomes purified from testis (Gomez *et al*., 2019; Xiong *et al*., 2022; Zhang *et al*., 2021; Zivkovic *et al*., 2022). As proteasomes in mature sperm nuclei are free of such factors, as shown by our cryo-ET data (Fig. 1), it will be important to investigate proteasome structures in differentiating germ cells to understand the nature of α4s-mediated interactions.

Previous studies reported that β1i, β2i and β5i immuno-subunits replace the canonical proteolytic β-subunits in the sperm proteasome (Qian *et al*., 2013). However, our sperm 20S proteasome structure did not contain immuno- or thymo-proteasome subunits (Fig. 3c; Extended Data Fig. 6c-f).

Remarkably, our high-resolution 20S structure shows an Arg-containing tripeptide covalently linked to the catalytic threonine of β2 subunits, a feature not previously reported to our knowledge (Fig. 4). There are several possible reasons for this. First, our gentler purification method, compared with previous studies (Harshbarger *et al*., 2015; Schrader J., 2016), preserves proteasomes in their active state throughout the process and may also supply substrate up to the cryo-EM sample preparation step (Extended Data Fig. 4 a-e). In particular, nuclease treatment at the beginning of the purification, combined with a reducing environment throughout the process, likely released monomeric protamines. As small peptides, these could enter the 20S CP for degradation, and remain visible in the active site due to low activity of the 20S CP (Coux, 1996). While the sperm 20S gate in our structure adopts a closed conformation, various studies indicate that it can be opened by oxidised and unfolded proteins with exposed hydrophobic patches (Grune, 2003; Jung *et al*., 2014; Latham *et al*., 2014; Orlowski, 2003) or by peptides containing the C-terminal HbYX (hydrophobic-tyrosine-any amino acid) motif (Opoku-Nsiah *et al*., 2022; Smith, 2007). Indeed, although protamines are highly positively charged proteins, they do include a version of the XbYX motif in their sequence (residues 15-18, Tyr-Tyr-Arg; UniProt ID: P04553). During proteolysis by the β2 subunit, the protamine degradation product containing the Tyr-Tyr-Arg motif at the C-terminus could be formed and released, opening the proteasomal gate. Similarly, an early study suggested that polycationic substances, including protamines, enhance proteasomal activity (Mellgren, 1990), and more recent data propose proteasomal activation by polyamines (Kudriaeva *et al*., 2020). This supports the idea that protamines could be degraded by the sperm 20S CP. The sperm proteasome gate consists of an N-terminal insertion in α4s* that positions a positively charged R4^α4s*^ at the inner surface, whereas the same region is neutral in the canonical variant (Extended data Fig. 7c). This may represent an adaptation for processing cationic protamine substrates, affecting both substrate access and the release of cleavage products. In mature spermatozoa, protamines are crosslinked by disulfide bridges, preventing their degradation (Balhorn, 2007). The process may instead be initiated post-fertilisation.

The peptide observed in the active site reveals a native substrate in the β2 subunit for the first time. Unexpectedly, neither Cys31^β2^, Gly45^β2^, nor His35^β2^ coordinates the substrate (Fig. 4c), distinguishing β2 from other catalytic subunits (Groll M., 1997; Harshbarger *et al*., 2015; Huber *et al*., 2016). This is likely due to the distal positioning of the positively charged group on the substrate side chain. In yeast, a Gly45A^β2^ mutation does not affect cell growth, subunit folding, or ligand binding, consistent with the proposal that Gly45^β2^ is not necessary for substrate recognition (Xin *et al*., 2019). The position of the water molecule previously proposed to act as a nucleophile resolving the acyl-enzyme intermediate (Groll M., 1997; Huber *et al*., 2016; Marques *et al*., 2009) is suboptimal for efficient nucleophilic attack. A similar conclusion was drawn in a study reporting the high-resolution structure of a somatic proteasome in complex with inhibitors (Schrader J., 2016). While Schrader *et al*. proposed that a water molecule hydrogen-bonded to the active site Tyr169 and Thr21 plays a critical, yet unknown role, the corresponding water molecule does not coordinate the peptide in our structure. Altogether, the mechanism of resolving the acyl-enzyme intermediate requires further study and may involve conformational changes, likely facilitated by engagement of proteasomal activators.

In our 20S-PA200 structure, the peptide density was nearly absent in the activator-proximal β2 subunits, suggesting that it represents a trapped catalytic intermediate in the 20S CP. Decreased substrate occupancy is accompanied by conformational changes in the activator-proximal β2 active site of 20S-PA200 structure (Fig. 4e). While previous work on the canonical 20S-PA200 structure (Toste Rego and da Fonseca, 2019) identified rearrangements of the trypsin site upon PA200 binding, the presence of a substrate in the active site revealed the mechanistic implications of these conformational changes. To explore this, we directly compared the effect of conformational changes in PA200-distal and -proximal β2 subunits on substrate binding, in the single map (Fig. 4d-e). The conformation-dependent differences in substrate processing rates proposed here may apply to canonical proteasomes as well, due to the conservation of the β2 subunit between sperm and somatic 20S-PA200 proteasomes. Previous studies have suggested that α4s may favour association with the PA200 activator (Gomez et al., 2019; Zhang et al., 2021). However, in our structure, the residues at the α4s–PA200 interface are largely conserved relative to those in the α4 subunit (Extended Data Fig. 6a–c; Extended Data Fig. 8a–d; Zivkovic et al., 2022). Our data, therefore, suggests that α4s* may not alter the intrinsic binding preferences of the 20S proteasome toward PA200, in agreement with Zivkovic et al. (Zivkovic *et al*., 2022).

Interestingly, in human sperm cells and tissue, 19S was enriched in the tail of differentiating spermatids and mature spermatozoa, an enrichment not shared by 20S subunits (Extended Data Fig. 9f-j). Several studies have proposed that α4s confers sperm 20S preference for 19S (Wang *et al*., 2025; Zivkovic *et al*., 2022). However, our structural data does not support this notion, as there are no interactions between α4/α4s and 19S in our map (Extended Data Fig. 8e-g). Therefore, it remains unclear how specific nuclear clustering between 20S and PA200, but not between 20S and 19S, occurs in the sperm head during differentiation. The lack of 19S complexes in the nucleus and their exclusive localisation at the human sperm tail may be due to the presence of additional factors, although further work is required to validate this hypothesis.

Our human primary tissue data place proteasomal cluster formation concurrent with protamination and hypercondensation of DNA in round spermatids (Extended Fig. 9k-l). At this stage, histones are replaced first with transition proteins and subsequently with protamines (Christensen, 1984; Rathke, 2014), a process that is dependent on proteasomal activity (Qian et al., 2013). In agreement with this, we show that post-meiotic nuclear proteasome clusters in human tissue co-localise with the PA200 activator (Fig. 5b-c), which confers specificity for histones targeted for removal through hyperacetylation (Christensen, 1984; Qian *et al*., 2013). The replacement of histones with protamines ultimately results in hyper-compaction of chromatin (Pogany, 1981). It is therefore possible that, during DNA compaction, proteasome complexes become excluded from the DNA, leading to the formation of *lacunae.* In our cryo-tomographic and STORM data of spermatozoa nuclei (Fig. 1, 2), proteasome-containing *lacunae* are indeed found deeply embedded between the highly condensed protamine-DNA phase. They may be retained there due to extreme DNA condensation, protamine crosslinking, and disrupted nucleocytoplasmic transport (Santos *et al*., 2024).

Various studies have proposed roles for the sperm proteasome during and after fertilisation (Rawe *et al*., 2008; Sutovsky *et al*., 2004; Wang *et al*., 2025). Our data did not reveal proteasome complexes in the acrosomal region; rather, proteasome clusters were found almost exclusively in the nucleus, with some also present in the tail. Further studies of capacitated sperm could help clarify what role sperm proteasomes may have in the degradation of the zona pellucida, as previously proposed (Sutovsky *et al*., 2004; Zimmerman *et al*., 2011). A recent report proposed that the sperm 20S proteasome binds oocyte-derived 19S regulatory particles to degrade Fetuin B and prevent polyspermy (Wang *et al*., 2025). However, 20S proteasomes have been shown to be present in oocytes (Assou *et al*., 2009; Wang *et al*., 2025; Zaffagnini *et al*., 2024; Zhang, 2025) and, according to our structural data, α4s* does not promote preferential binding to 19S. It is, however, plausible that α4s* interactors in the cytoplasm of the zygote drive proteasomal activity after fertilisation. Future structural work of zygote proteasomes is needed to clarify these pathways.

Finally, since protamine disulfide bonds are reduced in mature oocytes (Jenkins and Carrell, 2012; McLay, 2003), 20S sperm proteasomes from *lacunae* may be responsible for protamine degradation in the zygote, driving decondensation of the paternal genome (Assou *et al*., 2009; Wang *et al*., 2025; Zaffagnini *et al*., 2024; Zhang, 2025).

Overall, our integrated structural and cellular analyses reveal the *in situ* structure, organisation and compositional variance of the nuclear human sperm proteasome and uncover sperm-specific features. Our high-resolution structures provide molecular insights into substrate processing, the role of PA200 and of the testis-specific subunit α4s, which we find to be a specific splice variant in human sperm cells. Together, our study provides a structural framework for future studies on proteasome function and regulation during sperm-cell differentiation and early embryonic development.

## Methods

### Human sperm and testis tissue samples

Human mature sperm cells from 25 healthy donors were obtained from the European Sperm Bank. The samples had been processed by density gradient centrifugation prior to cryopreservation. Donors have been tested by the European Sperm Bank according to the requirements of the national competent authorities and EU Commission directive 2006/17/EC, before samples were released. Consent to use the samples for research purposes was obtained by the European Sperm Bank. Human sperm straws were thawed and used immediately. Before grid vitrification or immunostaining experiments on glass, sperm cells were checked for mobility using an optical microscope. Human testis FFPE sections (approximately 5 μm thick) from five healthy donors, between 18 and 55 years of age and with no history of infertility, were obtained from the Imperial College Healthcare Tissue Bank. All the experimental procedures were approved by the Imperial College Healthcare Tissue Bank (Research Ethics Committee approval numbers: 22/WA/0214; ICHTB HTA license: 12275).

Human testis FFPE sections (5 µm thick) from >10 individuals, aged 18-55 years of age and with no history of infertility, were obtained from the Imperial College Healthcare Tissue Bank (Department of Surgery and Cancer), the Human Research Tissue Bank (Cambridge University Hospitals) and commercially (AMSBIO, cat. HP-401, BP-401). Experimental procedures were performed under REC approval (REC Ref. 24/WM/0226, IRAS Project ID: 340922) and the Imperial College Healthcare Tissue bank (Research Ethics Committee approval numbers: 22/WA/0214; ICHTB HTA license: 12275).

### Mouse sperm and tissue

Mouse sperm from WT C57BL/6 JAX mice was used for this study. Mouse sperm straws were thawed, and cells were used immediately after checking for motility. Mouse (C57BL/6) testes FFPE sections (5 µm thick) were purchased from AMSBIO.

### Sample preparation and cryo-FIB milling

Quantifoil R 1/4 Au 200 mesh EM grids (Quantifoil Micro Tools GmbH) were glow discharged (Edwards S150B glow discharger) at 30 mA for 45 sec on both sides of the grid. A Leica GP2 or a custom-made manual plunger were used for plunge freezing in liquid ethane, after blotting for 9-12 seconds, using a sperm concentration of around 1×10^5^ – 5×10^5^ cells/grid. Vitrified cells were thinned to 150-200 nm lamellae using a Scios cryo-FIB/SEM (Thermo Fisher Scientific), an Aquilos II cryoFIB/SEM (Thermo Fisher Scientific) or a Crossbeam 550 cryo-FIB/SEM (Zeiss GmbH). Milling was performed as previously described (Dendooven *et al*., 2025; Santos *et al*., 2024). Briefly, inorganic platinum was sputtered on the grid in a Quorum chamber (Quorum Technologies) for the Scios and the Crossbeam 550, or in the main chamber for the Aquilos II. After that, the cells were coated with organometallic platinum (Trimethyl [(1,2,3,4,5-ETA.)-1 Methyl-2, 4-Cyclopentadien-1-YL] Platinum) using a gas injection system (GIS). A milling angle of 7-9° relative to the grid’s plane was used for milling, using a stepwise staircase pattern with the following currents (1nA, 0.5nA, 0.3nA, 0.1nA and 30pA).

### Cryo-ET data collection, tomogram reconstruction and subtomogram averaging

270 tilt-series of the sperm nucleus were collected on lamellae using Serial EM PACE-tomo (Eisenstein *et al*., 2023). Tilt series were acquired using a 300kV Titan Krios G3i Transmission Electron Microscope (Thermo Fisher Scientific) equipped with a Bioquantum energy filter (Gatan) and a K3 direct-electron detector (Gatan) or equipped with a Selectris energy filter and Falcon4i detector. A dose-symmetric tilt scheme was used with a total fluence of 100e/A^2,^ (3 e/Å^2^ per tilt). Defocus was in the −2 to −5 µm range. Processing of tilt series was carried out using Warp (Tegunov and Cramer, 2019). Tilt series alignment was executed with Aretomo (Zheng *et al*., 2022) and tomograms were reconstructed at ∼10 Å/pix through weigthed backprojection and filtered to 60Å for particle picking. Proteasome particles were and picked using crYOLO (Wagner *et al*., 2019). 46,831 particles were used for initial map generation and alignment with Dynamo (Castaño-Díez D, 2012). RELION 3.1 (Zivanov *et al*., 2020) was used for refining the initial poses, resulting in a 19.5 Å map. Extensive masked 3D classification in RELION resulted in three classes (20S with 43,128 particles; 20S-PA200 with 2,430 particles; PA200-20S-PA200 with 245 particles) whose poses were refined with M (poses, image warp and stage angles were refined) (Tegunov *et al*., 2021) to obtain a 20S map at 6.8 Å resolution; a 20S-PA200 map at 9.5 Å resolution; and a PA200-20S-PA200 map at 14.4 Å resolution.

### Cryo-SEM tomography

Cryo-FIB/SEM volume imaging was performed with a Crossbeam 550 cryo-FIB/SEM (Zeiss GmbH) (Schertel *et al*., 2013), equipped with a PP3010Z cryo stage and loading system (Quorum Technologies). A carpet of vitrified sperm cells on grid was milled at a stage tilt of - 4° with a current of 300 pA (30 kV) and a slice thickness of 16 nm. The pixel size for SEM imaging was 4.7 nm for human and 4.98 nm for mouse sperm, using an InLens detector at 2.3kV and 50pA with line averaging. Volumes were processed in Fiji as previously described (Schindelin *et al*., 2012). Briefly, contrast-limited adaptive histogram equalisation was used to enhance contrast (Zuiderveld, 1994), then images were aligned using the SIFT algorithm (Lowe, 2004). Horizontal and vertical curtain stripes were removed by stripe suppression using bandpass filtering. 9 human and 5 mouse sperm cells were segmented manually using Imod (Kremer, 1996).

### Proteasome purification from human sperm cells

All the steps were performed at 4 °C. 200 x 10^6^ IUI-ready human sperm cells (European Sperm Bank Aps, Denmark) were centrifuged at 300 × g for 15 min. The pellet was resuspended in buffer L (25 mM HEPES pH 7.4, 10 % [v/v] glycerol, 1 mM ATP, 1 mM DTT, 125 U mL^-1^ Benzonase nuclease (Merck KGaA, Germany), 1 mg mL^-1^ DNase I (Merck KGaA, Germany)) and incubated for 30 min. The cells were sonicated for 5 min, homogenised and incubated for 30 min. Next, the homogenate was centrifuged at 20,000 × g for 10 min, and the supernatant was concentrated using Amicon Ultra Centrifugal Filter 100 kDa MWCO (Merck KGaA, Germany). The concentrated sample was loaded on Superose 6 increase 3.2/300 (Cytiva) equilibrated in buffer A (25 mM HEPES pH 7.4, 10 % (v/v) glycerol, 100 mM NaCl, 1 mM ATP, 1 mM DTT, 5 mM MgCl_2_). The eluted fractions were assayed for proteasome presence using immunoblotting and negative-stain transmission electron microscopy (NS-EM). Selected fractions were concentrated using Amicon Ultra Centrifugal Filter 100 kDa MWCO and loaded on Superose 6 increase 3.2/300 (Cytiva) equilibrated in buffer B (25 mM HEPES pH 7.4, 100 mM NaCl, 1 mM ATP, 1 mM DTT, 5 mM MgCl_2_). The fractions were assayed for proteasome presence using immunoblotting and NS-EM, concentrated and immediately used for single-particle transmission electron cryo microscopy (cryo-EM) sample preparation.

### Immunoblotting

The protein fractions were mixed at a 1:1 ratio with 1x NUPAGE LDS sample buffer (Invitrogen) supplemented with 50 mM DTT and separated on NUPAGE 4-12% Bis-Tris SDS-PAGE gels (Invitrogen). Subsequently, the samples were transferred onto PVDF membrane (Bio-Rad) using Trans-Blot semi-dry transfer system (Bio-Rad) and immunoblotted using an iBind Flex system (Thermo Fisher Scientific) according to the manufacturer’s instructions. The primary antibody, 20S Proteasome β7 (H-3) (sc-365725, Santa Cruz Biotechnology), was used at 1:1000 dilution. The secondary antibody, goat anti-mouse IgG (HRP) (ab97023), was used at 1:2000 dilution. Immunoblots were imaged using Chemidoc MP device (Bio-Rad).

### Negative-stain transmission electron microscopy (NS-EM)

To assess proteasome presence in the chromatographic fractions using NS-EM, EM copper grids (Carbon Films on 200 Mesh Copper Grids, Agar Scientific) were glow-discharged in the Edwards S150B glow discharger for 30 s at 40 mA and 0.1 mbar. Next, 3 µL protein sample was deposited on the glow-discharged grid and incubated for 30 s, RT, followed by three wash steps in ddH_2_O and staining in 2% (w/v) uranyl acetate for 1 min, RT. The samples were screened using a 120 kV Tecnai Spirit transmission electron microscope (Thermo Fisher Scientific) equipped with Orius CCD camera (Gatan) at 21,000x magnification and pixel size 2.53 Å/px.

### Single particle cryo-EM sample preparation

Quantifoil R1.2/1.3 Cu300 holey carbon grids were cleaned with chloroform, acetone, and isopropanol as described in Passmore and Russo, 2016 (Passmore and Russo, 2016). A 2.1 nm-thick continuous carbon layer was evaporated on a mica sheet (Agar Scientific) using the Leica EM ACE600 carbon coater according to the manufacturer’s instructions. Next, the carbon layer was floated on water and transferred onto the cleaned grids. Subsequently, the continuous carbon-coated grids were glow-discharged in the Edwards S150B for 30 s at 40 mA and 0.1 mbar and placed in Vitrobot Mark IV (Thermo Fisher Scientific) semi-automated plunger equilibrated at 4 °C and 100% relative humidity. Three microliters of concentrated proteasome sample was applied to the grid, incubated for 30s and blotted with Watman-1 filter paper for 2s at −7 blot force. Finally, the grid was vitrified in liquid ethane maintained at 93K using a cryostat device (Russo *et al*., 2016) and stored in liquid nitrogen.

### Single particle cryo-EM data collection

Cryo-EM data was collected on a 300 kV Titan Krios G4 microscope equipped with SelectrisX energy filter (EF) and Falcon 4i post-EF direct electron detector (Thermo Fisher Scientific) using aberration-free image shift. The energy filter slit was set at 10 eV width, and the defocus varied between −1.0 µm and −2.0 µm. Zero-loss frames were collected at 105,000x magnification and corresponding calibrated pixel size of 0.926 Å/ px^-1^ in EER format with 40 e-Å^2^ total fluence over 3s exposure. Four data sets were acquired. Dataset 1 comprised 9,158 micrographs of the 20S proteasome fraction (Extended Data Fig. 4e). Datasets 2, 3, and 4 comprised 9,832, 26,881, and 13,500 micrographs, respectively, of the 20S-PA200 proteasome fraction (Extended Data Fig. 4e).

### Single particle cryo-EM data processing

Dataset 1 was pre-processed in RELION 4.0.1 (Zivanov *et al*., 2022). Datasets 2, 3, and 4 were pre-processed in RELION 5.0.1 (Burt *et al*., 2024). For each dataset, 918 hardware EER frames were grouped into 45 fractions, resulting in 0.87 e-/ Å^2^ dose per fraction, and motion-corrected using MotionCor2 in patch mode (Zheng *et al*., 2017). The contrast transfer function (CTF) parameters were estimated with CTFFIND-4.1 (Rohou, 2015). Twenty micrographs of various defoci from dataset 1 were manually picked and used to train the neural network of crYOLO 1.9.7 (Wagner *et al*., 2019). The trained model was used for automated particle picking on each entire dataset. The particle number and map resolution for all processing steps are shown in Extended data Fig. 4f. The particles were extracted in RELION, imported into cryoSPARC v4.4.1 (Punjani, 2017), and 2D classified into 100 classes to remove picking false positives. The selected particles were used to generate five (dataset 1) or six (datasets 2, 3, 4) initial models. In each dataset, one of the maps corresponded to the intact, isotropic 20S reconstruction, while other maps consisted of defective and over-represented particles. Heterogeneous refinement was used to assign particles corresponding to each map. The particles corresponding to the best class were selected and re-imported into RELION using pyem (Asarnow, 2019). Datasets 3 and 4 were combined into a single star file with two optics groups. Each particle subset was 3D auto-refined followed by two iterations of per-particle CTF estimation, aberration corrections (Zivanov *et al*., 2020) and 3D auto-refinement. The particles from datasets 3 and 4 were split into separate star files and subjected to Bayesian polishing (Zivanov *et al*., 2019). The polished particles were imported into cryoSPARC v4.4.1. Particles from dataset 1 were processed separately, while datasets 2, 3, and 4 were combined with optic group separation. The sets were homogeneously refined with C2 symmetry, optimisation of per-particle defocus, per-group CTF parameters, and correction of the Ewald Sphere with negative curvature (Wolf, 2006). Next, the particles of datasets 2, 3, and 4 were split into 69 groups according to the beam-image shift parameters to refine the beam tilt, trefoil, and spherical aberration parameters for each group separately, followed by per-particle CTF estimation and homogeneous refinement with parameters corresponding to the “Homogeneous refinement 1” step. The particles of datasets 2, 3, and 4 were subjected to 3D classification with 15 classes and a mask encompassing the region outside the 20S core particle. Class 1 corresponded to the 20S core particle bound to the PA200 activator (6,748 particles) while class 2 represented the 26S proteasome (4,207 particles). Each of the two classes was subjected to non-uniform reconstruction with default parameters, resulting in maps of 2.9 Å and 3.3 Å global resolution at Fourier Shell Correlation (FSC) of 0.143, for 20S-PA200 and 26S, respectively. The remaining particles from datasets 2, 3, and 4, as well as the particles from dataset 1 – Homogeneous refinement 1, were re-imported in RELION using pyem. A RELION script, eer_trajectory_handler.py, was used to change the rendering mode to the 8K grid and group each 8 EER frames into a single fraction. Each particle set was subjected to Bayesian polishing with particle re-extraction at a super-resolution pixel size of 0.668 Å px^-1^ and re-imported into cryoSPARC. The particles from each dataset were split into 69 groups according to the beam-image shift parameters. Following homogeneous refinement, with C2 symmetry, minimisation over per-particle scale, optimisation of per-particle defocus and per-group CTF parameters (tilt, trefoil, spherical aberration, tetrafoil, anisotropic magnification), as well as Ewald sphere correction with a negative curvature, the particles were 3D classified into 10 classes at 5Å filtering resolution, with initial class similarity of 0.8 and the “auto-tune initial class similarity” option turned on. This revealed a single main class of 222,000 particles that was used for non-uniform refinement with C2 symmetry and Ewald sphere correction. The refinement resulted in a 1.85 Å resolution map, at the FSC threshold of 0.143.

### Model building

The asymmetric unit of the somatic 20S proteasome structure (PDB ID: 6RGQ) (Toste Rego and da Fonseca, 2019) was rigid-body fitted into the 1.85 Å map of the sperm 20S core particle in UCSF ChimeraX 1.9 (Pettersen *et al*., 2021). The α4s initial model was generated with AlphaFold2 (Jumper *et al*., 2021) using the sequence of the alternatively spliced variant of α4s (UniProt ID: Q8TAA3-5). The α4 subunit was replaced with α4s initial model in Coot 0.9 (Emsley and Cowtan, 2004) and refined in real-space with PHENIX 1.21.1 (Afonine *et al*., 2018). Water molecules surrounding the asymmetric unit were added to the map and validated using the ‘Check/delete waters’ tool in Coot. Following several iterations of manual rebuilding in Coot and real-space refinement in PHENIX 2.0-5884, the NCS operators were applied to generate a structure with C2 symmetry. Finally, the chains were renamed to match the 20S-PA200 structure in Coot. To confirm the presence of the alternatively-spliced α4s variant (UniProt ID: Q8TAA3-5), referred as α4s*, rather than the canonical one (UniProt ID: Q8TAA3-1, referred to as α4s FL), the asymetric unit of the somatic 20S proteasome structure (PDB ID: 6RGQ) (Toste Rego and da Fonseca, 2019) was rigid-body fitted into the 1.85 Å map of the sperm 20S core particle in UCSF ChimeraX 1.9 (Pettersen *et al*., 2021). The α4 subunit was replaced with α4s AlphaFold2 model (UniProt ID: AF-Q8TAA3-F1) in Coot 0.9 (Emsley and Cowtan, 2004), manually rebuilt, and real-space refined in PHENIX 1.21.1 (Afonine *et al*., 2018). To generate the 20S-PA200 structure, the somatic 20S-PA200 coordinates (PDB ID: 6KWY) (Guan *et al*., 2020; Toste Rego and da Fonseca, 2019) were rigid-body fitted into the 2.9 Å map of the sperm 20S-PA200 proteasome with UCSF Chimera 1.9. The α4 subunit was replaced with α4s* coordinates from sperm 20S CP with Coot 0.9, followed by multiple iterations of real-space refinement in PHENIX 2.0-5884 and manual rebuilding in Coot. The models were validated with MolProbity (Williams *et al*., 2018). The figures were generated in UCSF ChimeraX 1.9.

### Immunofluorescence

Human and mouse sperm cells were seeded on #1.5 glass coverslips or ibidi 8-well chamber slides (GmbH), pre-functionalised with Poly-L-lysine (PLL), and allowed to dry, at RT for 5-10 minutes. Cells were then fixed in 4% PFA in PBS, followed by washes in PBS. Samples were quenched with 50mM NH_4_Cl in PBS for 15 minutes at RT, followed by washes in PBS. Cells were permeabilised and simultaneously blocked in 3% bovine serum albumin (BSA), 0.2% Triton-X100 in PBS, for 1 hour at RT. Following this, cells were incubated for 2h at RT, or overnight at 4°C, with primary antibodies, diluted in 3% BSA, 0.2% Triton X-100 in PBS. Cells were washed with 0.5% BSA, 0.05% Triton X-100 in PBS and incubated for 1 hour at RT with fluorescently labelled secondary antibodies diluted in 3% BSA, 0.2% Triton X-100 in PBS. Following this, cells were washed in 0.5% BSA, 0.05% Triton X-100 in PBS, followed by PBS and mounted using Prolong Gold (Thermo Fisher Scientific) for SIM experiments or left in PBS (ibidi chambers) for STORM experiments.

### Stochastic Optical Reconstruction Microscopy and Cluster Analysis

Immunofluorescently labelled cells were imaged using a Nanoimager (ONI) system, with a 100x 1.49 NA (TIRF) oil immersion objective. Imaging was performed using 561 nm and 640 nm wavelength illumination, with acquisition of 15,000 frames (50,000 for DNA) and 30 ms exposure time. Acquisition was performed using Highly Inclined and Laminated Optical Sheet (HILO) illumination and laser power 60-150 mW. Imaging was performed using ONI Bcubed buffer (ONI). For two-colour STORM measurements, a calibration procedure with TetraSpeck beads (Invitrogen) was used for channel alignment. STORM data was processed using CODI software (ONI), including drift-correction and HDBSCAN analysis. For H(Ram *et al*., 2010) analysis, a minimum cluster size was set at 15 molecules.

### Structured Illumination Microscopy

Structured Illumination Microscopy (SIM) was performed using a Zeiss Elyra S.1 system, equipped with a 63x 1.46NA plan-apochromat oil immersion objective; 405 nm, 488 nm, 561 nm and 647 nm laser lines and a CMOS camera. 0.091 µm Z-stacks were collected with 3 or 5 rotations. For two-colour imaging, channel alignment was performed with mounted TetraSpeck beads (Invitrogen), using the affine method. Data analysis was performed using Zeiss Zen Black 2.3 (Zeiss).

### Immunohistochemistry

Pre-mounted FFPE-embedded testis sections (5 µm thick) from human healthy donors or C57BL/6 mice (5 µm thick) were deparaffinised and rehydrated. Tissue sections were then subjected to heat-induced antigen retrieval, submerged in 10 mM Sodium Citrate, 0.05% Tween 20 (pH 6). Tissue sections were incubated in 0.05% Triton-X100 in TBS for 15 min with rocking at room temperature, followed by 30 min incubation in blocking solution (1% BSA, 10% Donkey Serum in TBS) at RT. Sections were then incubated overnight at 4°C with primary antibodies diluted in 1% BSA in TBS. The next day, tissue sections were washed 3 times in 0.05% Triton X-100 in TBS, for 10 min at room temperature, with rocking. Tissue sections were then incubated for 1 h at RT with fluorescently labelled secondary antibodies diluted in 1% BSA in TBS. Tissue was then stained with Hoechst 33342 (Thermo Fisher Scientific) in 0.05% Triton-X100 in TBS for 15 min at RT. This was followed by three 10 min washes in 0.05% Triton-X100 in TBS, at RT, a TBS wash and a final wash in diH2O. Samples were mounted using #1.5 coverslips and Prolong Gold Mounting media (Thermo Fisher Scientific).

### Confocal Microscopy

Tissue sections were imaged using an Andor Bench Top Confocal BC43 (Oxford Instruments), equipped with a 60x 1.42NA oil immersion objective, 405 nm, 488 nm, 561 nm and 638 nm lasers. Fusion software (Oxford Instruments) was used for image acquisition. Image analysis was performed using Fiji (ImageJ).

## Data availability

Subtomogram averages of the 20S, PA200 and PA200-20S-PA200 were deposited in the Electron Microscopy Data Bank (EMDB) with accession codes EMD-55396, EMD-55397, and EMD-55398, respectively. Single particle cryo-EM maps for 20S, 20S-PA200, and 26S were deposited in EMDB with accession codes EMD-56072, EMD-56074, and EMD-56075, respectively. The atomic coordinates of 20S and 20S-PA200 were deposited in the Protein Data Bank (PDB) with accession codes PDB ID: 9TMN and 9TMS, respectively. All structural data is available to reviewers.

## Acknowledgements

We thank the Allegretti and Barford lab members for useful discussions and critical feedback. We thank David Barford for reading part of the manuscript and providing valuable insights on structure interpretation. We thank Dom Bellini for advice on atomic modelling. We thank Mart Last for support with tomogram denoising. We thank David Haselbach and Jan Löwe for scientific discussions on proteasomes. We thank Emmanuel Derivery’s lab for the acetylated tubulin antibody. We thank the cryo-EM Facility at LMB for maintaining the electron microscopes, the Scientific Computing facility for maintaining our computer cluster and the Light Microscopy Facility for maintaining light microscopes. We thank the Frederic Langevin from the LMB transgenic facilities and the LMB Ares animal facility for providing mouse sperm cells used in this study. We acknowledge Diamond for access and support at the Aquilos-II of the UK national electron Bio-Imaging Centre (eBIC). For the purpose of open access, the MRC Laboratory of Molecular Biology has applied a CC BY public copyright licence to any Author Accepted Manuscript version arising. M.A., T.D., P.K., were funded by the UKRI Medical Research Council (MC_UP_1201/30), A.d.S., and O.K. were funded by the Wellcome Trust Early Career Award to A.d.S. (227622/Z/23/Z).

## Author Contributions

P.K., A.d.S., T.D., and M.A. conceived and designed the experiments; T.D. and A.d.S. collected cryo-FIB milling, cryo-SAV and cryo-ET data; T.D., A.d.S., and M.A. analysed cryo-ET data; T.D. performed subtomogram averaging; M.A. analysed cryo-SAV data; A.d.S. performed immunohistochemistry, SIM, STORM, and confocal microscopy, and analysed all the data; O.K. performed immunofluorescence; P.K. and A.d.S performed protein purification; P.K. performed single-particle cryo-EM acquisition and analysis; A.d.S., T.D. and M.A. supervised the study. P.K., A.d.S., T.D., and M.A. wrote the manuscript.

## Ethics declarations

Human mature sperm cells were purchased from the European Sperm Bank (https://www.europeanspermbank.com/en). The samples had been previously processed, tested by the bank according to the requirement of the national competent authorities, GMP and WHO standards, as well as EU directives: 2006/86/EC, 2006/17/EC, 2004/23/EC, 2015/565. Sperm cells were screened for morphology, genetic diseases, blood-borne viruses and motility. Consent to use the samples for research purposes was given to the European Sperm Bank. Human testis FFPE sections from five different individuals, aged 18-55 years of age and with no history of infertility, were obtained from the Imperial College Healthcare Tissue Bank (Department of Surgery and Cancer). All the experimental procedures were approved by the Imperial College Healthcare Tissue bank (Research Ethics Committee approval numbers: 22/WA/0214; ICHTB HTA license: 12275).

## Material & correspondence

Material requests and correspondence should be addressed to Matteo Allegretti

---

**Extended Data Video 1. The molecular architecture of a large proteasome-rich nuclear *lacuna*.** Summarising movie displaying the three proteasome maps from sub-tomogram averaging mapped back in a denoised cryo-tomogram of the sperm nucleus.

**Extended Data Video 2. Volumetric cryo-EM of human sperm cells shows nuclear *lacunae*.** Movie displaying cryo slice-and-view imaging of several sperm cells with nuclear lacunae segmented in green in the central cell. Scale bar = 1 µm.

**Extended Data Table 1.** Statistics of cryo-EM data collection, data processing, and model refinement.

**Extended Data Fig. 1.**
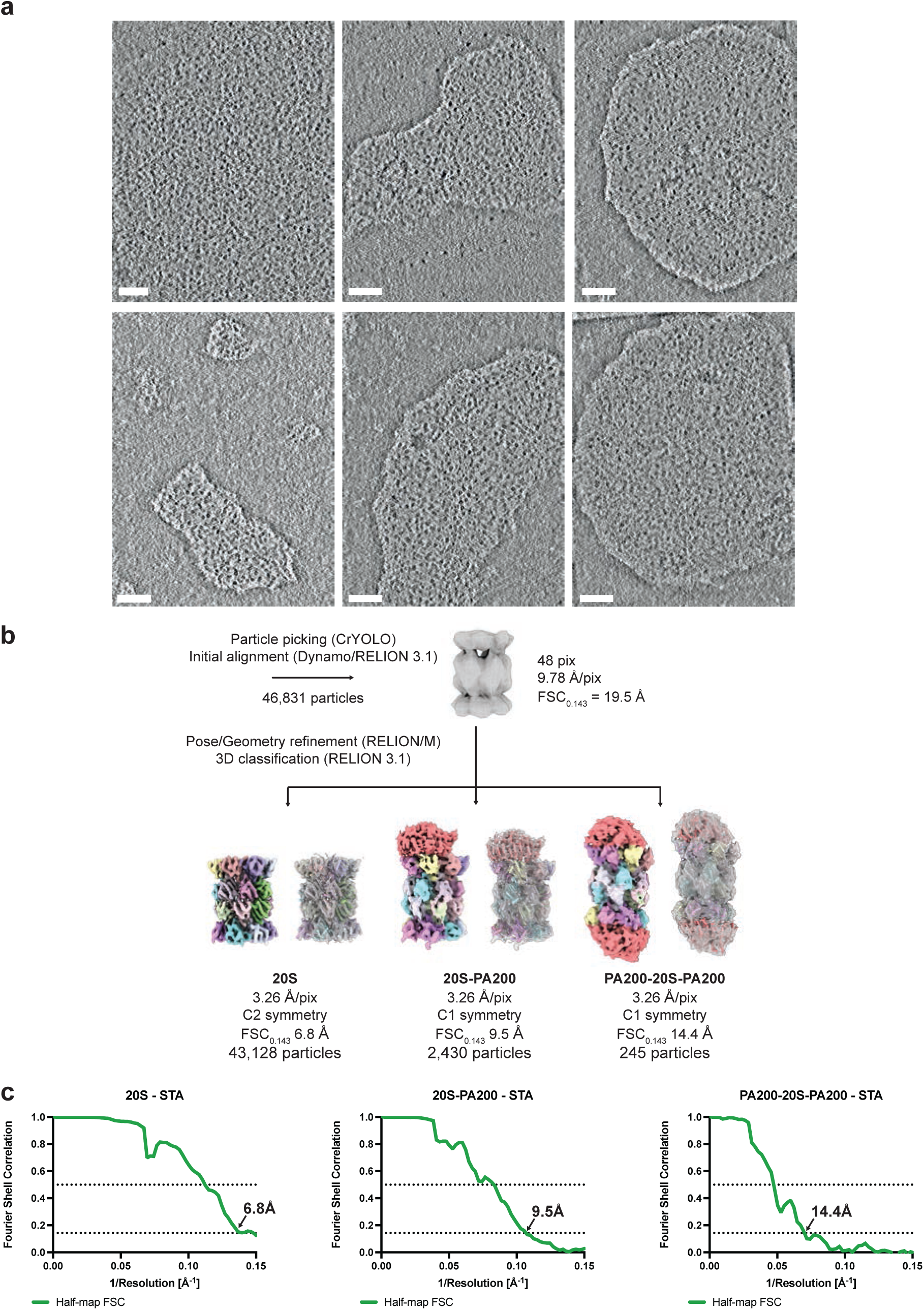
Gallery of lacunae and subtomogram averaging processing pipeline of nuclear sperm proteasomes. **a,** cryo-tomographic slices from deconvolved tomograms of six nuclear lacunae filled with barrel-shaped proteins. Scale bar = 100 nm. **b,** Details of the subtomogram averaging processing pipeline of 20S, 20S-PA200 and PA200-20S-PA200 proteasomes. Subtomogram averages with fitted PDBs are shown. **c**, Gold standard Fourier Shell Correlation (FSC) curves for the 20S (which reached Nyquist), 20S-PA200 and PA200-20S-PA200 proteasome reconstructions (STA stands for subtomogram averaging). The resolution values at FSC = 0.143 (half-map FSC) are indicated.

**Extended Data Fig. 2.**
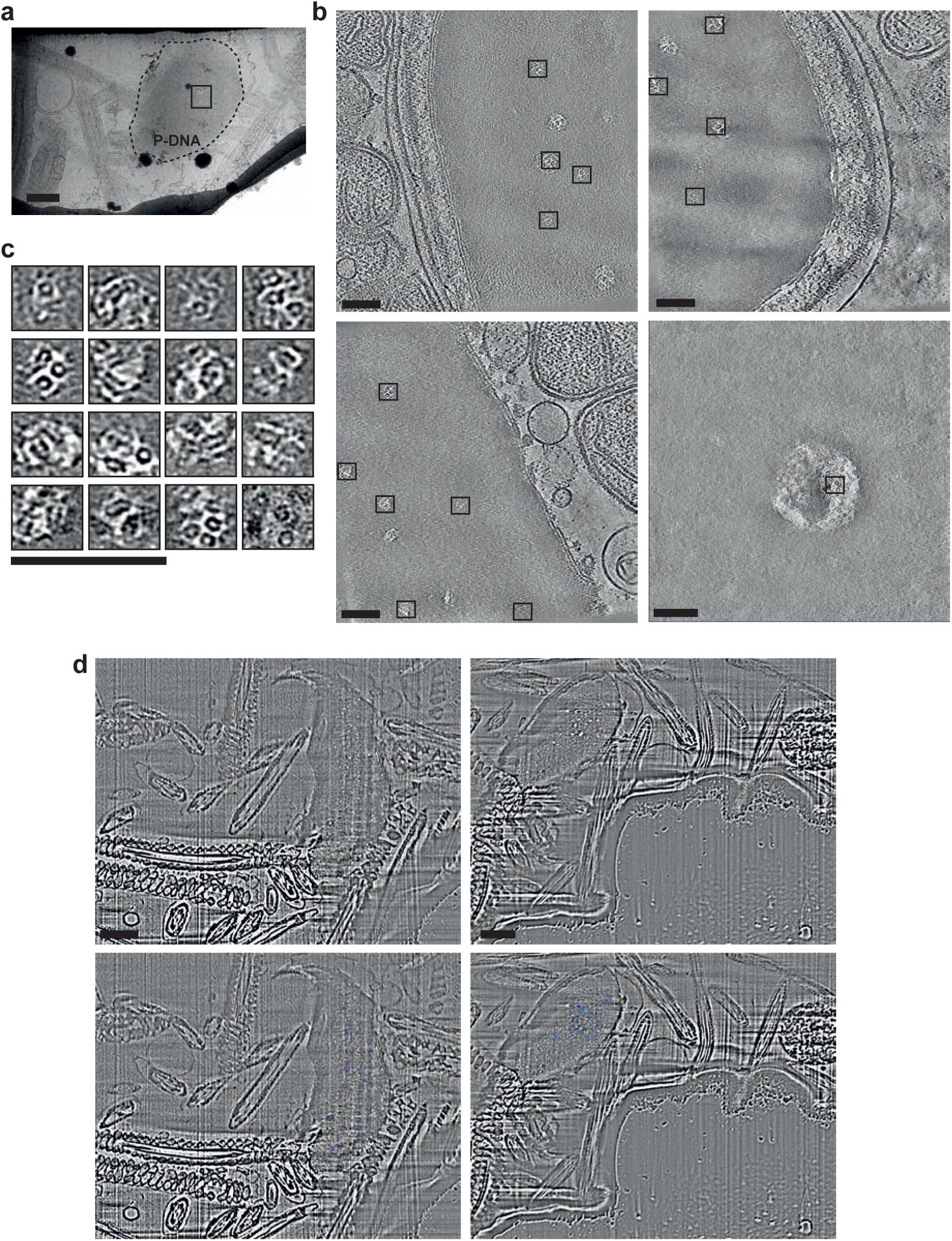
Proteasome complexes in mouse sperm nuclear *lacunae.* **a,** cryo-TEM low magnification lamellae map showing one mouse sperm head. Arrows indicate regions with sperm *lacunae* targeted for tilt-series acquisition. Scale bar = 1 *μ*m. **b,** z-slices from reconstructed tomograms showing barrel-shaped protein complexes inside membrane-less nuclear compartments. Scale bar = 100nm. **c,** zoom-in regions from tomograms shown in **b**, with barrel-shaped proteins inside *lacunae*. Scale bar = 100nm. **d,** representative z-slices of 3D reconstructed cryo-SEM volumetric data of mouse sperm cells. Small sperm *lacunae* are visible within the nucleus. Scale = 1 *μ*m. Bottom panels show detail of segmented *lacunae* in colour (purple and blue).

**Extended Data Fig. 3.**
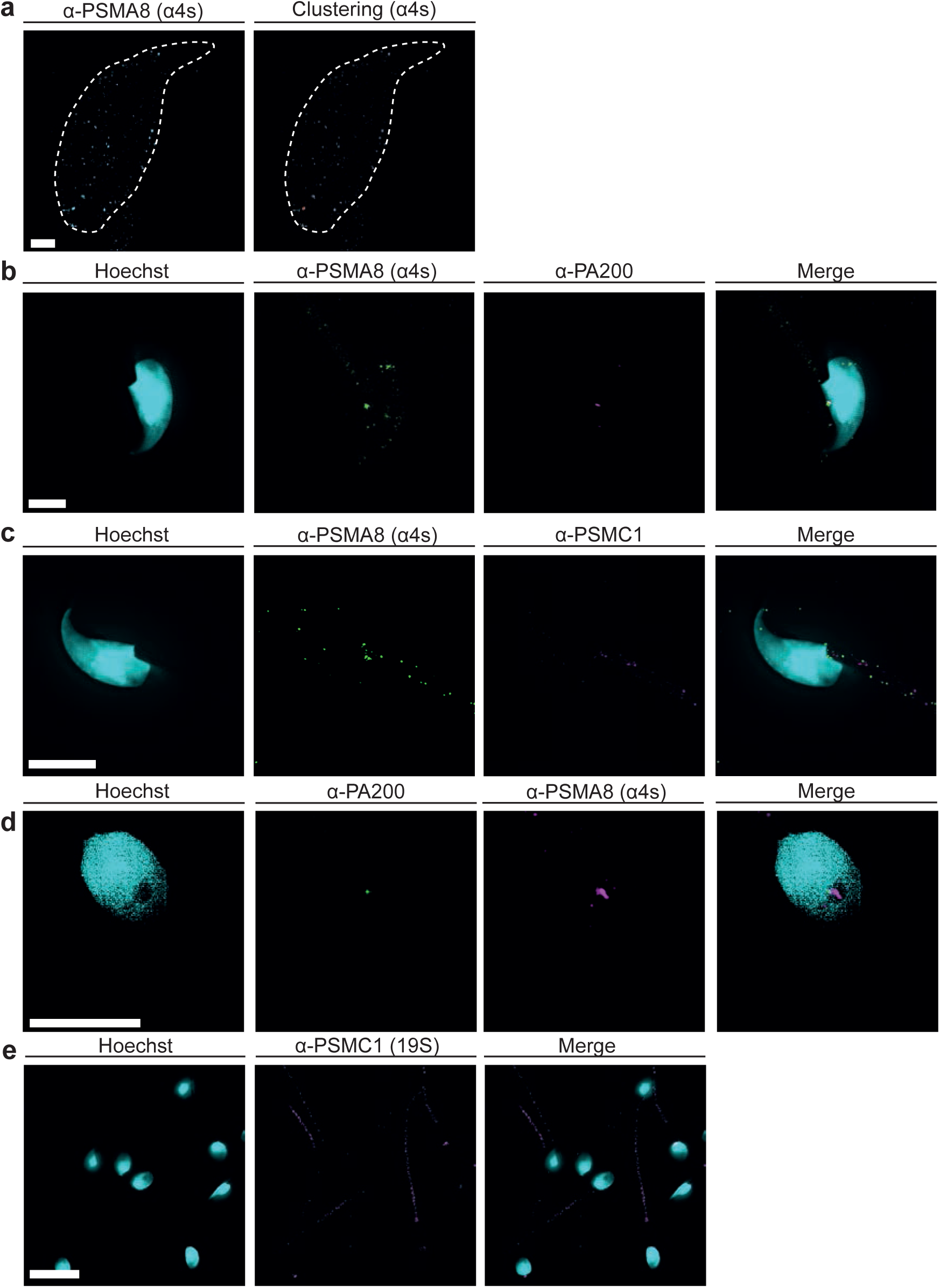
Proteasome distribution and clustering in mouse and human sperm cells. **a,** STORM of sperm-specific proteasome subunit PSMA8 (***α***4s) in a mouse sperm cell. The white dashed line shows the sperm nuclear region. The right panel shows ***α***4s cluster identification in red, superimposed to non-clustered localisations (cyan/grey) following DBSCAN analysis. Scale bar = 1 *μ*m. **b,** SIM image of mouse sperm cell showing foci of ***α***4s (green) and partial co-localisation with less abundant PA200 (magenta). Scale bar = 5 *μ*m. **c,** SIM image of mouse sperm cell showing localisation of ***α***4s and 19S subunit PSMC1. Scale bar = 5 *μ*m. **c,** SIM image showing localisation of ***α***4s in the nucleus of human sperm cells, segregated from DNA signal (Hoechst, cyan). PA200 signal can be observed partially localising to the same region of the nucleus. Scale bar = 5 *μ*m. **d,** SIM image showing localisation of 19S subunit, PSMC1 (magenta), in the tail of human sperm cells. Scale bar = 10 *μ*m.

**Extended Data Fig. 4.**
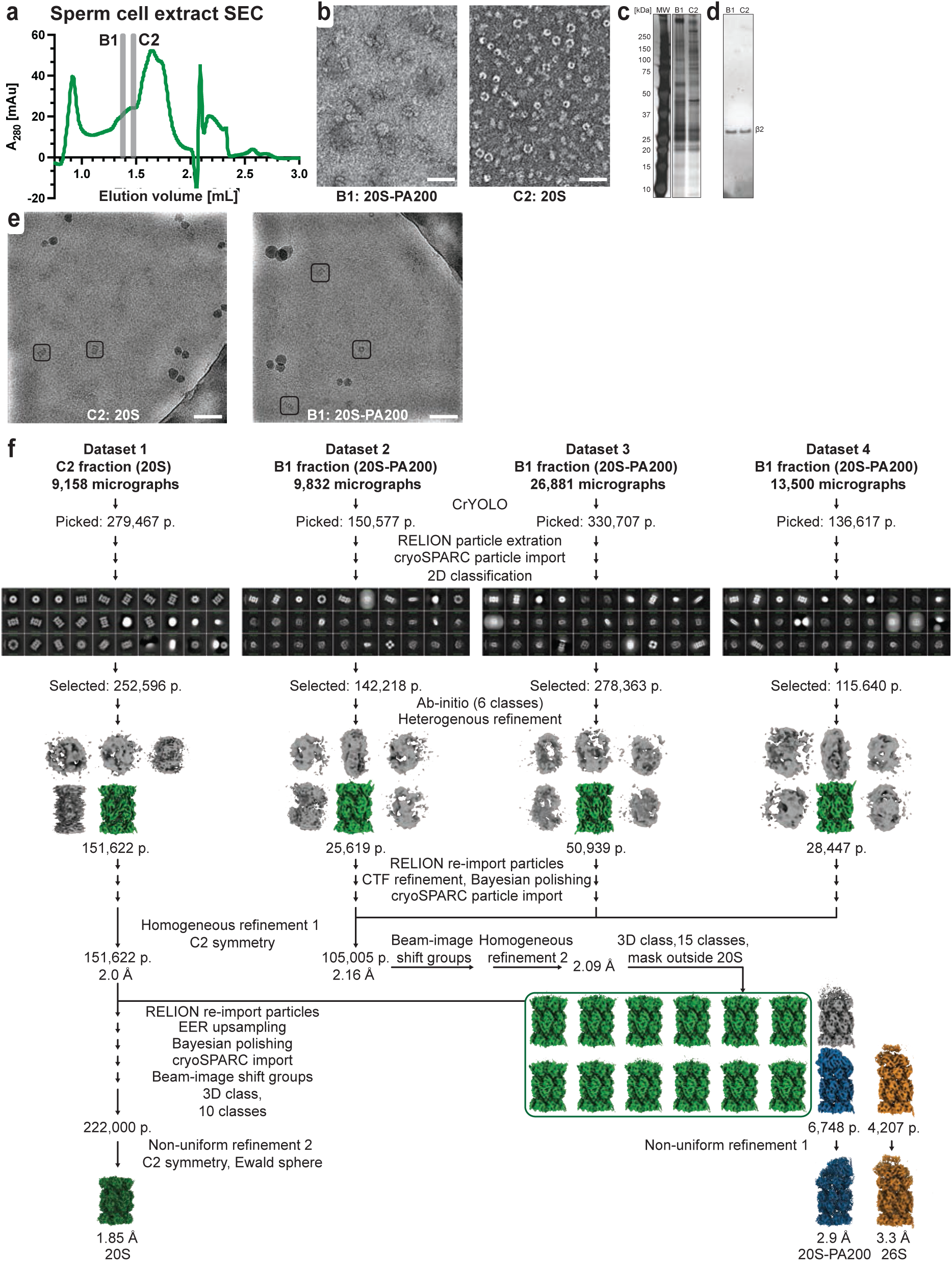
Purification and single-particle cryo-EM data processing of human sperm proteasomes. **a,** size exclusion chromatography (SEC) of the sperm cell extract. **b**, negative-stain transmission electron microscopy micrographs corresponding to SEC B1 and C2 fractions from **a**. **c,** silver-stained SDS-PAGE of B1 and C2 fractions. **d,** immunoblotting confirms the proteasome presence in B1 and C2 fractions. **e,** cryo-EM micrographs of the C2 (left) and B1 (right) fractions. Protein particles are marked with a black box. In **b** and **e**, the scale bars correspond to 50 nm. **f,** single-particle cryo-EM data processing pipeline. “p.” - particle number at corresponding processing step. 2D class averages of the initial particle selection, as well as maps from key processing steps, are shown.

**Extended Data Fig. 5.**
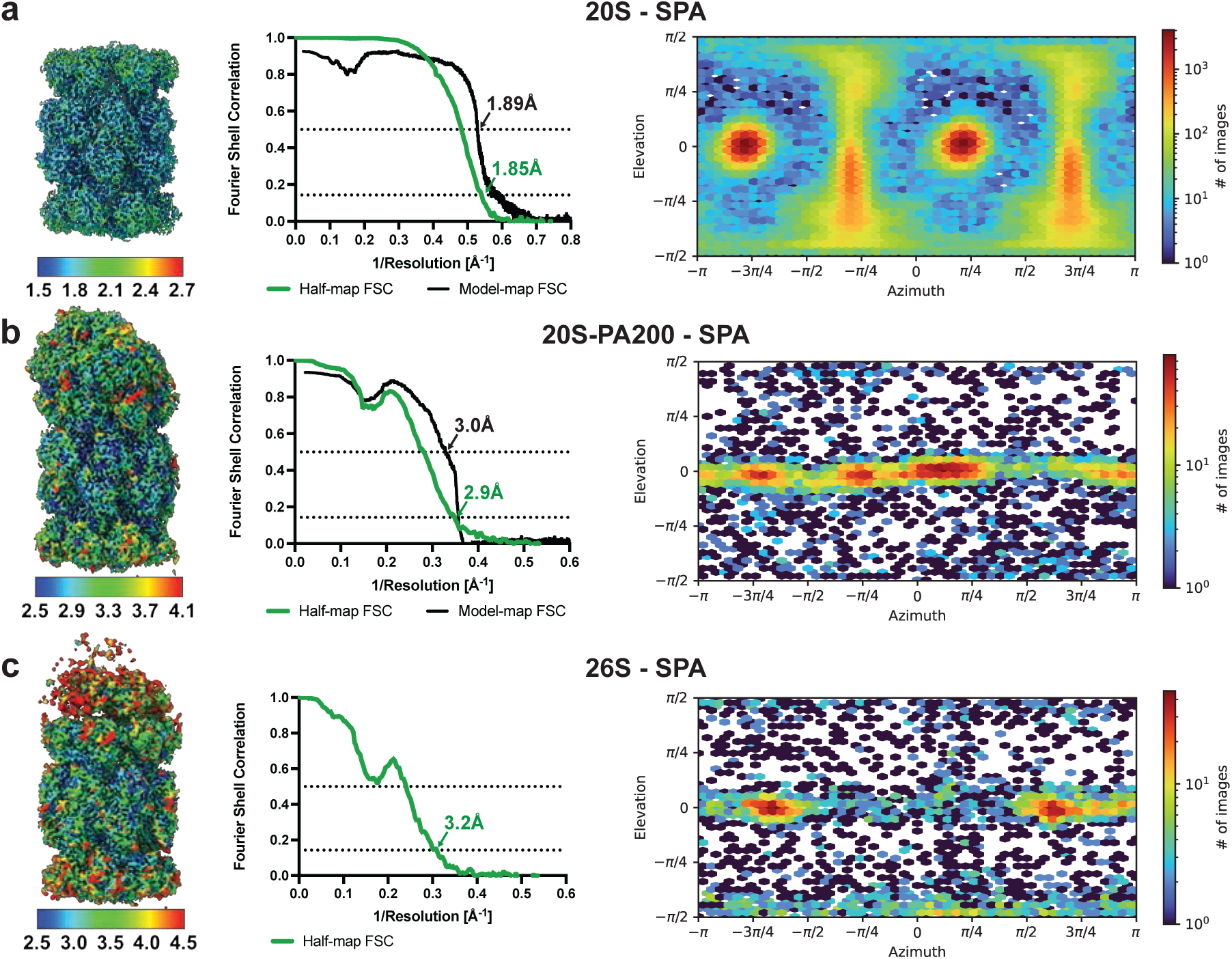
Properties of the final single-particle cryo-EM reconstructions (SPA). Local resolution maps (left), Fourier Shell Correlation (FSC) plots (middle), and angular distribution maps (right) are shown for the a) 20S, b) 20S-PA200, and c) 26S maps. In **a** and **b**, both the gold-standard half-map FSC (green) and the model-map FSC (black) are shown. The resolution values at FSC = 0.143 (half-map FSC) and FSC = 0.5 (model-map FSC) are indicated. The maps are locally coloured and filtered according to the resolutions of the colour key under each map.

**Extended Data Fig. 6.**
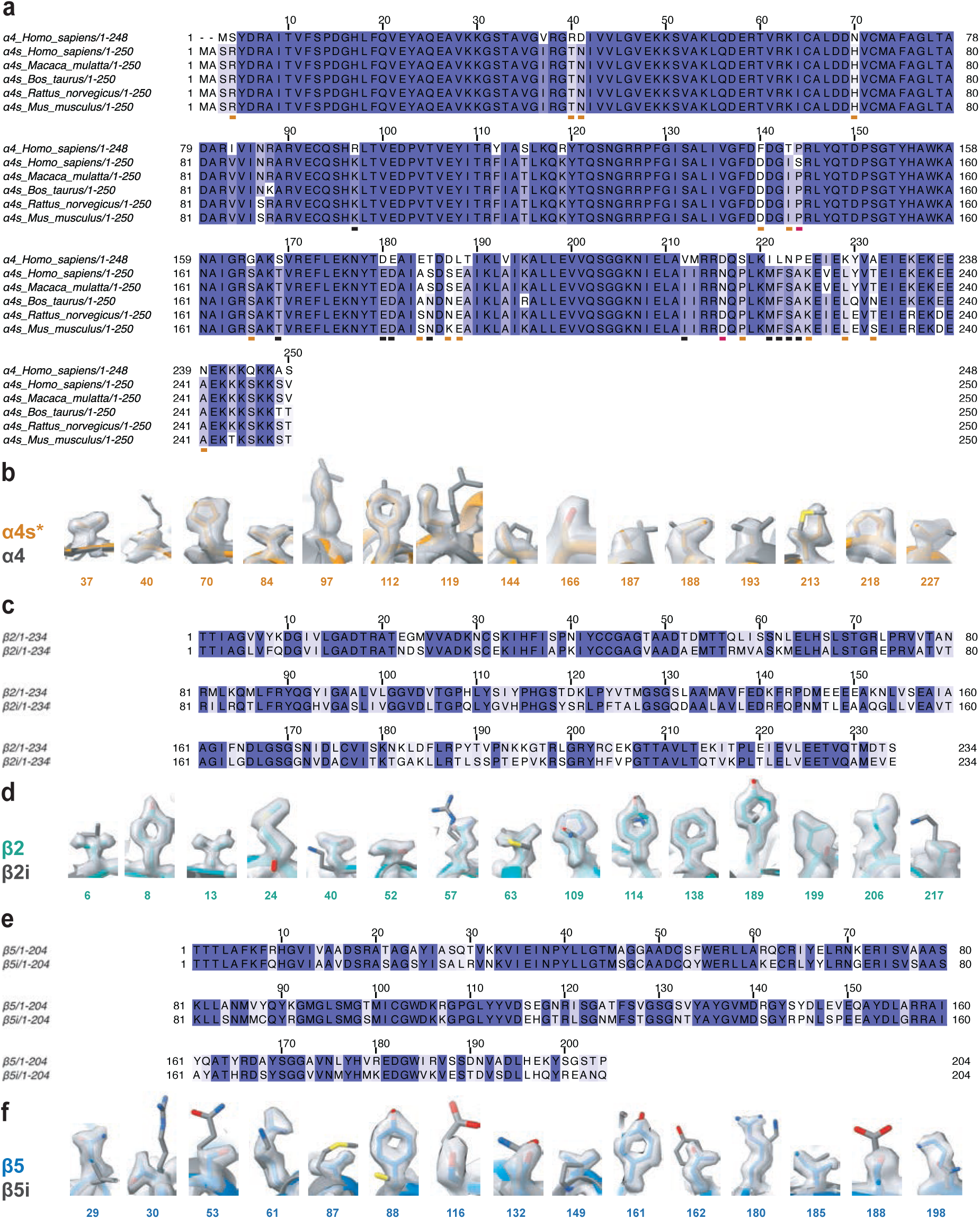
Subunit composition of the sperm proteasome. **a,** alignment of *Homo sapiens* α4 sequence (UniProt ID: O14818) with α4s sequences from five different species: *Homo sapiens* (α4s*, UniProt ID: Q8TAA3-5), *Macaca mulatta* (UniProt ID: A0A1D5Q503), *Bos taurus* (UniProt ID: A0AAA9S8Q9), *Rattus norvegicus* (UniProt ID: F1M6I7), *Mus musculus* (UniProt ID: Q9CWH6). The sequences are coloured by conservation. The underlines indicate solvent-exposed residues and are coloured by side-chain group conservation between α4 with α4s: black – conserved, orange – different side-chain group across all species, red – different side-chain group in some species. **b,** side-chain densities in the cryo-EM 20S proteasome map with modelled α4 (grey, PDB ID: 5LE5) and α4s* (yellow) side chains. The α4s* residue numbers are indicated. **c, e,** Sequence alignments of β2 (Uniprot ID: Q99436) and β2i (UniProt ID: P40306) **(c)** as well as β5 (UniProt ID: P28074) and β5i (UniProt ID: P28062) (**e)**, coloured by conservation. The propeptide sequences are omitted. **d, f,** side-chain densities in the cryo-EM 20S proteasome map with modelled side chains of β2 (cyan) and β2i (grey, PDB ID: 7b12) **(d)**, as well as β5 (blue) and β5i (grey, PDB ID: 7b12) **(f)**. The residue numbers are indicated.

**Extended Data Fig. 7.**
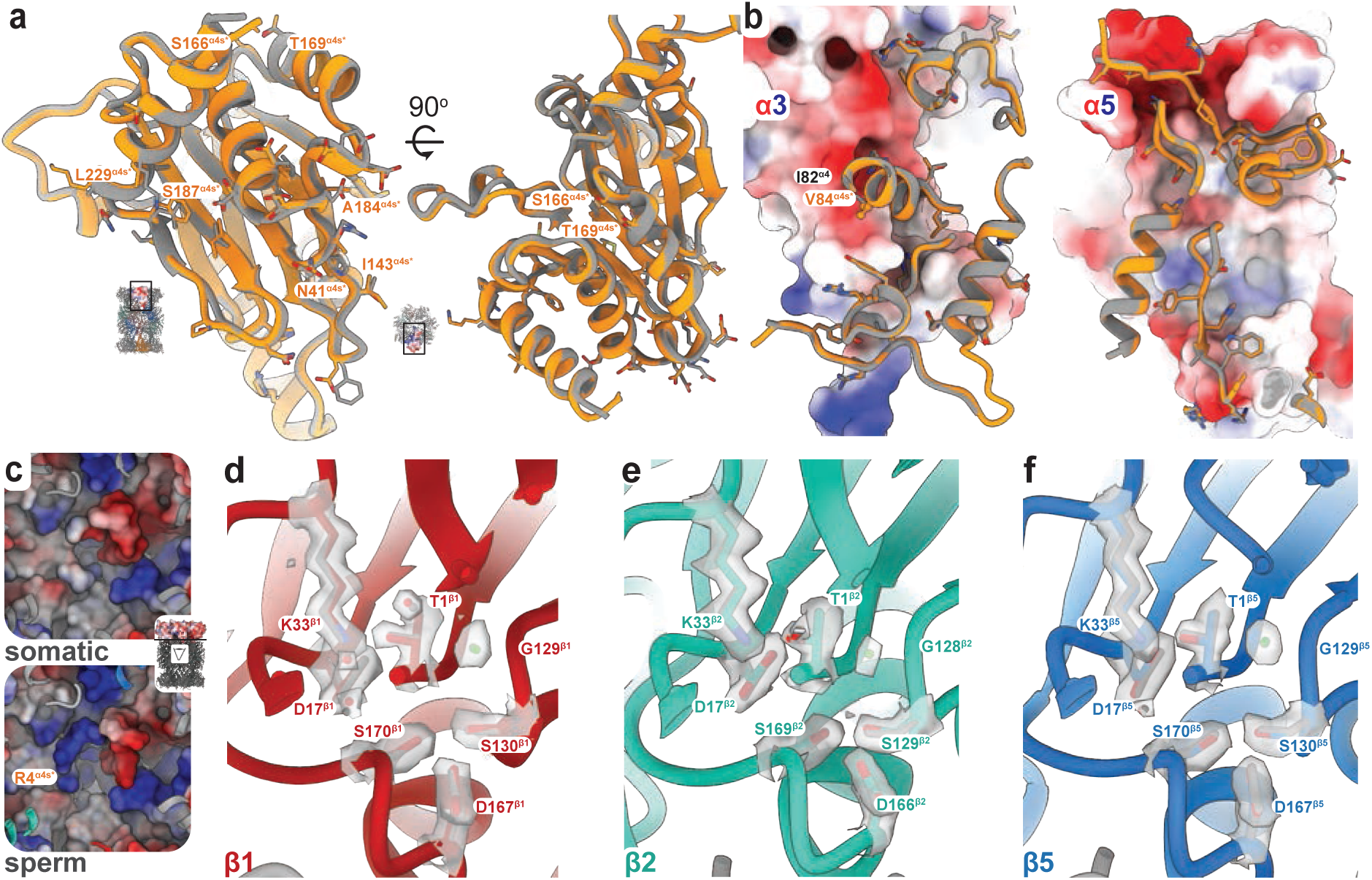
Structural features of the 20S sperm proteasome. **a,** α4s* structure (yellow) is highly conserved with α4 (grey, PDB ID: 5LE5). The models are shown in cartoon representation, and all non-conserved residues are shown as sticks. The solvent-exposed, non-conserved residues with a different side-chain group are indicated, including the substitutions creating the uncharged patch on the surface pf α4s*: Asp39^α4^ → Asn41^α4s^, Thr141^α4^ → Ile143^α4s^, Glu182^α4^ → Ala184^α4s^, Asp185^α4^ → Ser187^α4s^, and Lys227^α4^ → Leu229^α4s^ **b,** interfaces between α4s* (cartoon, yellow) or α4 (cartoon, grey) and α3 (left, shown as electrostatic surface) or α5 (right, shown as electrostatic surface) subunits. All interfacial residues of α4s*/α4 are shown as sticks, and the Ile82^α4^ → Val84^α4S^ mutation is indicated. **c,** the pores of somatic (PDB ID: 5LE5) and sperm 20S proteasome, shown as electrostatic surfaces and viewed from the catalytic chamber side, as indicated in the inset. R4^α4S*^ is indicated. **d, e, f** sperm 20S proteasome active sites in β1 (red, **d**), β2 (cyan, **e**), and β5 (blue, **f**) subunits, shown as cartoons. Catalytic residues are shown as sticks, along with their corresponding densities. The conserved catalytic water molecule is in green. The corresponding residues in the somatic proteasome (PDB ID:5LE5) are shown as grey sticks. In all panels, amino acids are marked using single-letter amino acid code.

**Extended Data Fig. 8.**
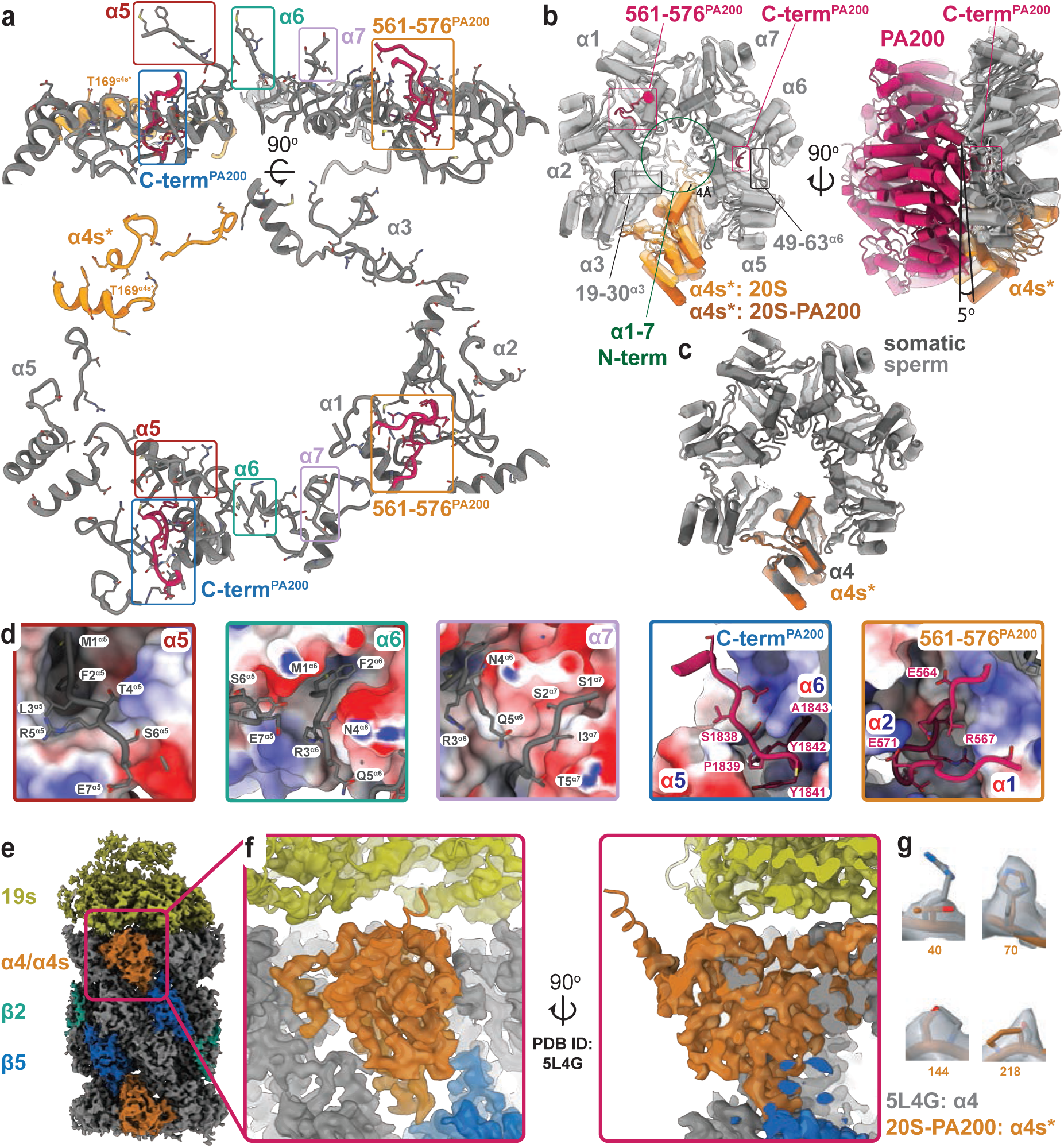
Interactions between the sperm 20S core particle and proteasomal activators. **a-d,** binding PA200 to sperm 20S CP opens the proteasomal gate. To achieve this, the 561-576^PA200^ loop extends into the α1-α2 interface while the C-terminal 1838-1843^PA200^ (C-term^PA200^) stretch displaces the α5-α6 subunits away from each other and shifts the 49-63^α6^ loop downwards. Channel opening is further facilitated by disordering the gate-forming, N-terminal loops of α1-α4s (α1-7 N-term), the flexibility of the first α3 helix (residues 19-30^α3^), and incorporating the α5, α6, and α7 subunit N-termini into the pockets on the PA200 surface. **a,** all the interactions between the α-subunits and PA200 activator in sperm 20S-PA200 structure. The α-subunit residues interacting with PA200 are shown as sticks. The secondary structural features harbouring these residues are shown as cartoons. The subunit names, as well as the only PA200-interacting residue in α4s* different to α4, Thr169^α4s^ replacing Ser167^α4^, are indicated. The rectangles mark structural features detailed in **d**. The main PA200 features interacting with the CP, i.e. the PA200 C-terminus and the 561-576 loop are shown as purple cartoons and sticks. **b,** conformational changes in the α-subunits occurring upon PA200 engagement lead to a 4 Å horizontal shift and a 5° downwards tilt of the α4s*. The structures are shown as cartoons with α-helices as tubes. The 20S-PA200 structure is opaque in darker tones, while the 20S is transparent and in brighter tones. The α4s is coloured in orange tones. The key features undergoing conformational changes, as well as the shift and tilt of α4s*, are indicated. **c,** comparison of somatic (PDB ID:6KWY, darker tones) and sperm (brighter tones) 20S-PA200 structures at the α-ring. α4 (grey) and α4s* (dark orange) subunits are indicated. RMSD between Cαs of all α-subunits is 0.599 (Guan *et al*., 2020). **d,** the main features stabilising the 20S-PA200 interface. The panels depict regions marked by rectangles in (**a**). In α5, α6, and α7 panels, PA200 is shown as an electrostatic surface, while the 20S is shown as cartoons and sticks. In C-term^PA200^ and 561-576^PA200^ panels, the 20S subunits are shown as electrostatic surfaces, with names indicated, while PA200 is shown as purple cartoons and sticks. **e,** cryo-EM map of the 26S proteasome isolated from human sperm. The 19S activator, α4/α4s*, β2 and β5 subunits are annotated. **f,** the interface between 19S and α4/α4s*. The 26S proteasome structure (PSB ID:5L4G) was rigid-body fitted in the map and shown as cartoons. **g,** side-chain densities in the sperm cryo-EM 26S proteasome map with modelled α4 (grey, PDB ID: 5L4G) and α4s* (orange, 20S-PA200 coordinates) side chains. In all panels, amino acids are marked using single-letter amino acid code.

**Extended Data Fig. 9.**
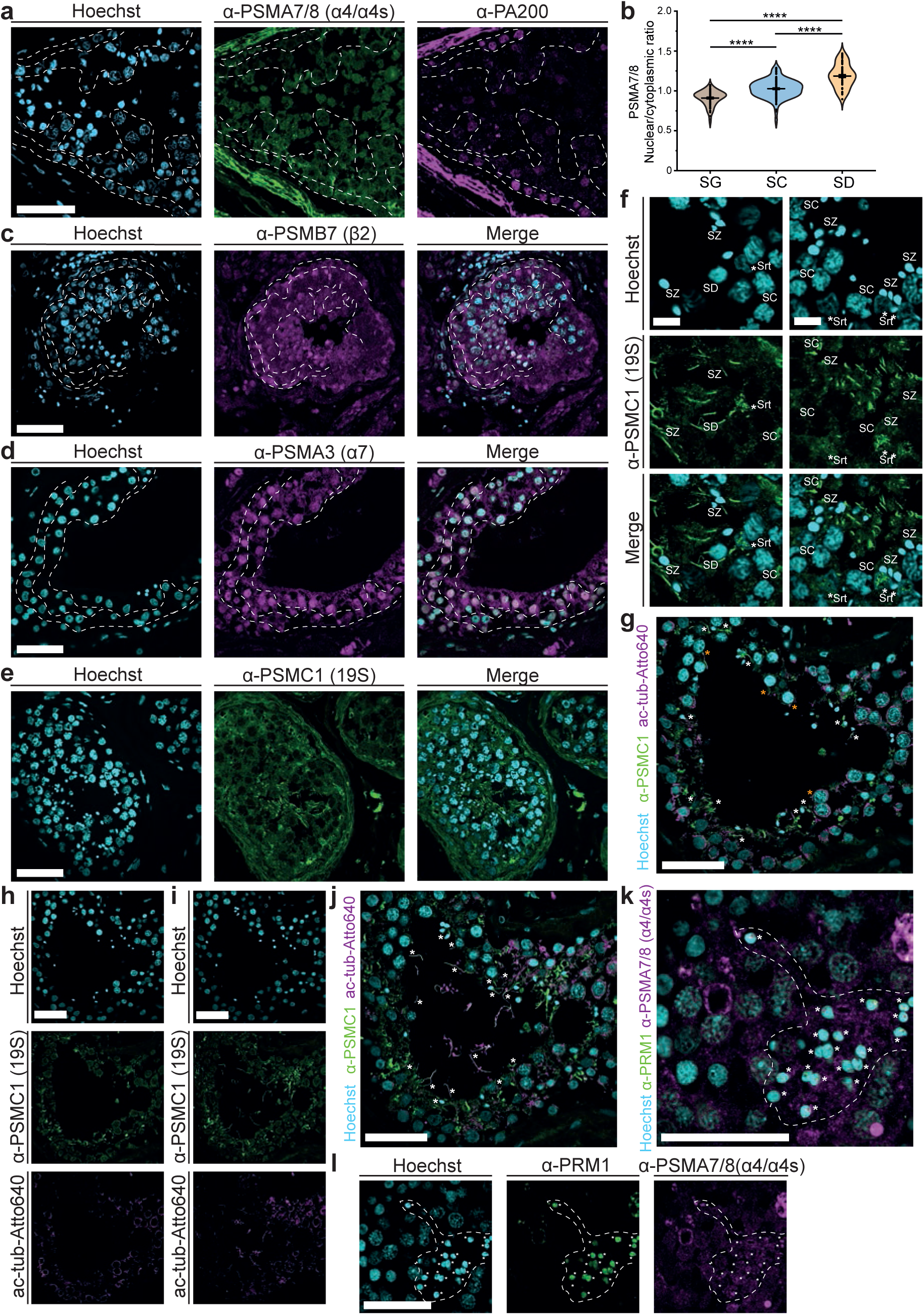
Distribution of proteasomal subunits and regulatory caps PA200 and 19S in human tissue during spermatogenesis. **a,** single-channel confocal image for merge shown in Fig. 5b, showing an overview of a seminiferous tubule from a healthy human donor. Immunohistochemistry staining shows distribution of proteasome subunits PSMA7/8 (*α*4/*α*4s) in green, proteasome adaptor PA200 in magenta and DNA (Hoechst, in cyan). White dashed lines represent boundaries between different cell types, as shown in Fig. 5a. Scale bar = 50 *μ*m. **b**, Quantification of nuclear/cytoplasmic proteasome ratios in differentiating germ cells. Quantification was performed from immunohistochemistry confocal microscopy data. Mean ± s.e.m. values are as follows: SG = 0.91 ± 0.01 (n= 83 cells, tissue from 3 donors); SC = 1.03 ± 0.01 (n= 136 cells, tissue from 3 donors); SD = 1.19 ± 0.02 (n=70 cells, tissue from 3 donors). Statistical analysis: One-way ANOVA Bonferroni-corrected (*\*\*\*\*p*< 0.0001). *p* = 2.6×10^-11^ (SG-SC); *p* = 1.9×10^-35^ (SG-SD); *p* = 2.9×10^-17^ (SC-SD). **c,** Distribution of PSMB7 (β2) subunit in human testis (magenta). DNA (Hoechst) is shown in cyan. Dashed lines show regions of different spermatogenic cell types. Scale bar = 50 *μ*m. **d,** Distribution of PSMA3 (*α*7) subunit in human testis (magenta). DNA (Hoechst) shown in cyan. Dashed lines show regions of different spermatogenic cell types. Scale bar = 50 *μ*m. **e,** Distribution of PSMC1, a subunit of the regulatory cap 19S in human testis (green). DNA (Hoechst) shown in cyan. Scale bar = 50 *μ*m. **f,** localisation of 19S subunit PSMC1 (green) to the mature tail of SZ, growing tail of SD and nucleus of Sertoli cells (*Srt). Scale bar = 10 *μ*m. **g,** merge showing co-localisation of PSMC1 (19S subunit, in green) with acetylated tubulin (magenta). Putative SC cilia are highlighted with orange stars. Other regions of co-localisation, including SZ and SD tails, are highlighted with white stars. DNA is shown in cyan (Hoechst). Scale bar = 50 *μ*m. **h,** individual channels for merge shown in *g***. i,** additional example of PSMC1 (19S, green) localisation relative to acetylated tubulin (magenta) in a human seminiferous tubule. Scale bar = 50 *μ*m. **j,** merge of images shown in **h.** Regions of co-localisation are highlighted with white stars. Scale bar = 50 *μ*m. **k,** confocal image merge showing clustering of proteasomes (PSMA7/8, magenta) in the nucleus of differentiating cells in human testes, relative to protamine signal (PRM1, green). Spermatids and spermatozoa are shown within the white-dashed line. Scale bar = 50 *μ*m. **l,** single-channel images for merge shown in k. Scale bar = 50 *μ*m.

**Extended Data Fig. 10.**
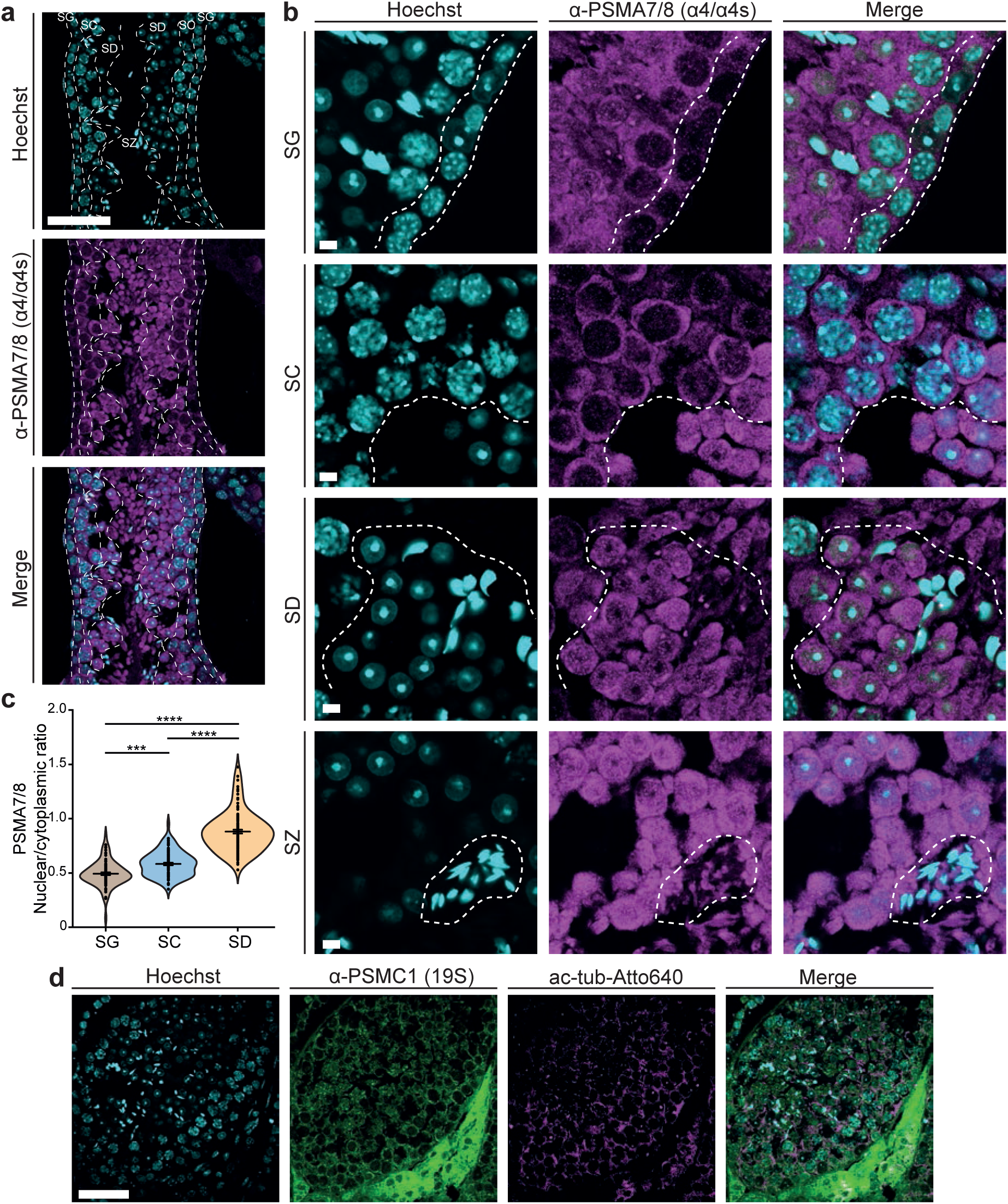
Distribution of proteasomal subunits and regulatory cap 19S during mouse spermatogenesis. **a,** confocal microscopy image showing mouse seminiferous tubule and distribution of proteasomal PSMA7/8 (*α*4/ *α*4s) subunits during sperm cell differentiation. White-dashed lines show tissue sub-compartments enriched for different germ cells (SG, SC, SD and SZ). Scale bar = 50 *μ*m. **b,** zoom-in region of mouse seminiferous tubules showing proteasome subunit PSMA7/8 (*α*4/ *α*4s) distribution during differentiation. White-dashed lines show regions with the different cell types: SG, SC, SD and SZ. Scale bar = 10 *μ*m. **c,** quantification, from confocal immunohistochemistry mouse data, of nuclear/cytoplasmic proteasome ratios in germ cells. Mean ± s.e.m. values are as follows: SG = 0.49 ± 0.01 (n= 80 cells); SC = 0.58 ± 0.01 (n= 88 cells); SD = 0.88 ± 0.02 (n= 112 cells). Statistical analysis: One-way ANOVA Bonferroni-corrected (*\*\*\*p*<0.001; *\*\*\*\*p*< 0.0001). *p* = 2.7 x10^-4^ (SG-SC); *p* = 2.3 x10^-48^ (SG-SD); *p* = 1.7 x10^-34^ (SC-SD). **d,** distribution of PSMC1 subunit of regulatory cap 19S (green) in mouse sperm testis, relative to acetylated tubulin (magenta). Scale bar = 50 *μ*m.

## References

Adolf, F., Du, J., Goodall, E. A.,…Schulman, B.A. (2024). Visualizing chaperone-mediated multistep assembly of the human 20S proteasome. Nat Struct Mol Biol 31, 1176–1188.

Afonine, P. V., Poon, B. K., Read, R. J.,…Adams, P.D. (2018). Real-space refinement in PHENIX for cryo-EM and crystallography. Acta Crystallogr D Struct Biol 74, 531–544.

Albert, S., Schaffer, M., Beck, F.,…Engel, B.D. (2017). Proteasomes tether to two distinct sites at the nuclear pore complex. Proc Natl Acad Sci U S A 114, 13726–13731.

Albert, S., Wietrzynski, W., Lee, C. W.,…Engel, B.D. (2020). Direct visualization of degradation microcompartments at the ER membrane. Proc Natl Acad Sci U S A 117, 1069–1080.

Allen, F. H. W., D. G.; Brammer, L.; Orpen, A. G.; Taylor, R. (2006). Chapter 9.5 Typical interatomic distances: organic compounds. Mathematical, physical and chemical tables C

Asarnow, D., Palovcak, E., Cheng, Y. (2019). UCSF pyem v0.5. Zenodo.

Assou, S., Cerecedo, D., Tondeur, S.,…De Vos, J. (2009). A gene expression signature shared by human mature oocytes and embryonic stem cells. BMC Genomics 10, 10.

Balhorn, R. (2007). The protamine family of sperm nuclear proteins. Genome Biol 8, 227.

Barad, B. A., Echols, N., Wang, R. Y.,…Fraser, J.S. (2015). EMRinger: side chain-directed model and map validation for 3D cryo-electron microscopy. Nat Methods 12, 943–6.

Burt, A., Toader, B., Warshamanage, R.,…Scheres, S. H.W. (2024). An image processing pipeline for electron cryo-tomography in RELION-5. FEBS Open Bio 14, 1788–1804.

Castaño-Díez D, K. M., Arheit M, Stahlberg H. (2012). Dynamo: a flexible, user-friendly development tool for subtomogram averaging of cryo-EM data in high-performance computing environments. J Struct Biol 178

Chemes, H. E. and Alvarez Sedo, C. (2012). Tales of the tail and sperm head aches: changing concepts on the prognostic significance of sperm pathologies affecting the head, neck and tail. Asian J Androl 14, 14–23.

Christensen, M. E. R., J. B.; Dixon, G. H. (1984). Hyperacetylation of histone H4 promotes chromatin decondensation prior to histone replacement by protamlnes during spermatogenesis in rainbow trout. Nucleic Acids Research 12

Coux, O., Tanaka, K., Goldberg, A. L. (1996). Structure and functions of the 20S and 26S proteasomes. Annu Rev Biochem 65

da Fonseca, P. C., He, J. and Morris, E. P. (2012). Molecular model of the human 26S proteasome. Mol Cell 46, 54–66.

Dendooven, T., Ebrahimi, M., dos Santos, A.,…Allegretti, M. (2025). BioRxiv

Dong, Y., Zhang, S., Wu, Z.,…Mao, Y. (2019). Cryo-EM structures and dynamics of substrate-engaged human 26S proteasome. Nature 565, 49–55.

Eisenstein, F., Yanagisawa, H., Kashihara, H.,…Danev, R. (2023). Parallel cryo electron tomography on in situ lamellae. Nat Methods 20, 131–138.

Emsley, P. and Cowtan, K. (2004). Coot: model-building tools for molecular graphics. Acta Crystallogr D Biol Crystallogr 60, 2126–32.

Enenkel, C., Kang, R. W., Wilfling, F. and Ernst, O. P. (2022). Intracellular localization of the proteasome in response to stress conditions. J Biol Chem 298, 102083.

Fleming, I. (2010). Ionic Reactions—Stereochemistry, Chapter 5. Molecular Orbitals and Organic Chemical Reactions

Gomez, H. L., Felipe-Medina, N., Condezo, Y. B.,…Pendas, A.M. (2019). The PSMA8 subunit of the spermatoproteasome is essential for proper meiotic exit and mouse fertility. PLoS Genet 15, e1008316.

Groll M., D. L., Löwe J., Stock D., Bochtler M., Bartunik H. D., Huber R. (1997). Structure of 20S proteasome from yeast at 2.4 A resolution. Nature 386

Grune, T., Merker, K., Sandig, G., Davies, K. J. (2003). Selective degradation of oxidatively modified protein substrates by the proteasome. Biochem Biophys Res Commun 305

Guan, H., Wang, Y., Yu, T.,…Ouyang, S. (2020). Cryo-EM structures of the human PA200 and PA200-20S complex reveal regulation of proteasome gate opening and two PA200 apertures. PLoS Biol 18, e3000654.

Guo, Q., Lehmer, C., Martinez-Sanchez, A.,…Fernandez-Busnadiego, R. (2018). In Situ Structure of Neuronal C9orf72 Poly-GA Aggregates Reveals Proteasome Recruitment. Cell 172, 696–705 e12.

Guo, X. (2022). Localized Proteasomal Degradation: From the Nucleus to Cell Periphery. Biomolecules 12

Hackerova, L., Klusackova, B., Zigo, M.,…Simonik, O. (2023). Modulatory effect of MG-132 proteasomal inhibition on boar sperm motility during in vitro capacitation. Front Vet Sci 10, 1116891.

Haraguchi, C. M., Mabuchi, T., Hirata, S.,…Yokota, S. (2007). Possible function of caudal nuclear pocket: degradation of nucleoproteins by ubiquitin-proteasome system in rat spermatids and human sperm. J Histochem Cytochem 55, 585–95.

Harshbarger, W., Miller, C., Diedrich, C. and Sacchettini, J. (2015). Crystal structure of the human 20S proteasome in complex with carfilzomib. Structure 23, 418–24.

Huber, E. M., Heinemeyer, W., Li, X.,…Groll, M. (2016). A unified mechanism for proteolysis and autocatalytic activation in the 20S proteasome. Nat Commun 7, 10900.

Jenkins, T. G. and Carrell, D. T. (2012). Dynamic alterations in the paternal epigenetic landscape following fertilization. Front Genet 3, 143.

Jiang, T. X., Ma, S., Han, X.,…Qiu, X.B. (2021). Proteasome activator PA200 maintains stability of histone marks during transcription and aging. Theranostics 11, 1458–1472.

Jumper, J., Evans, R., Pritzel, A.,…Hassabis, D. (2021). Highly accurate protein structure prediction with AlphaFold. Nature 596, 583–589.

Jung, T., Hohn, A. and Grune, T. (2014). The proteasome and the degradation of oxidized proteins: Part II - protein oxidation and proteasomal degradation. Redox Biol 2, 99–104.

Khor, B., Bredemeyer, A. L., Huang, C. Y.,…Sleckman, B.P. (2006). Proteasome activator PA200 is required for normal spermatogenesis. Mol Cell Biol 26, 2999–3007.

Klein, M., Busch, M., Friese-Hamim, M.,…Schadt, O. (2021). Structure-Based Optimization and Discovery of M3258, a Specific Inhibitor of the Immunoproteasome Subunit LMP7 (beta5i). J Med Chem 64, 10230–10245.

Kong, M., Diaz, E. S. and Morales, P. (2009). Participation of the human sperm proteasome in the capacitation process and its regulation by protein kinase A and tyrosine kinase. Biol Reprod 80, 1026–35.

Kremer, J. R. M., D. N.; McIntosh, J. R. (1996). Computer visualisation of three-dimensional image data using IMOD. J Struct Biol 116

Kudriaeva, A. A., Saratov, G. A., Kaminskaya, A. N.,…Belogurov, A. A., Jr. (2020). Polyamines Counteract Carbonate-Driven Proteasome Stalling in Alkaline Conditions. Biomolecules 10

Lafarga M, B. M., Pena E, Mayo I, Castaño JG, Bohmann D, Rodrigues JP, Tavanez JP, Carmo-Fonseca M. (2002). Clastosome: A Subtype of Nuclear Body Enriched in 19S and 20S Proteasomes, Ubiquitin, and Protein Substrates of Proteasome. Molecular Biology of the Cell 13

Latham, M. P., Sekhar, A. and Kay, L. E. (2014). Understanding the mechanism of proteasome 20S core particle gating. Proc Natl Acad Sci U S A 111, 5532–7.

Lee, J., Le, L., Kim, E. and Lee, M. J. (2021). Formation of Non-Nucleoplasmic Proteasome Foci during the Late Stage of Hyperosmotic Stress. Cells 10

Lowe, D. G. (2004). Distinctive Image Features from Scale-Invariant Keypoints. International Journal of Computer Vision

Marques, A. J., Palanimurugan, R., Matias, A. C., Ramos, P. C. and Dohmen, R. J. (2009). CatalyticMechanismandAssemblyof theProteasome. ChemRev 109

McLay, D. W., Clarke, H. J. (2003). Remodelling the paternal chromatin at fertilization in mammals. Reproduction 125

Mellgren, R. L. (1990). Interaction of human erythrocyte multicatalytic proteinase with polycations. Biochim Biophys Acta

Morales, P., Kong, M., Pizarro, E. and Pasten, C. (2003). Participation of the sperm proteasome in human fertilization. Hum Reprod 18, 1010–7.

Morales, P., Pizarro, E., Kong, M. and Jara, M. (2004). Extracellular localization of proteasomes in human sperm. Mol Reprod Dev 68, 115–24.

Opoku-Nsiah, K. A., de la Pena, A. H., Williams, S. K.,…Gestwicki, J.E. (2022). The YPhi motif defines the structure-activity relationships of human 20S proteasome activators. Nat Commun 13, 1226.

Orlowski, M., Wilk, S. (2003). Ubiquitin-independent proteolytic functions of the proteasome. Arch Biochem Biophys 1

Passmore, L. A. and Russo, C. J. (2016). Specimen Preparation for High-Resolution Cryo-EM. Methods Enzymol 579, 51–86.

Pérez-Moreno, I., López-Jiménez, P., Zapata, H.,…Gómez, R. (2025). The dynamics of ciliogenesis in prepubertal mouse meiosis reveal new clues about testicular maturation during puberty. BioRxiv

Pettersen, E. F., Goddard, T. D., Huang, C. C.,…Ferrin, T.E. (2021). UCSF ChimeraX: Structure visualization for researchers, educators, and developers. Protein Sci 30, 70–82.

Pogany, G. C. C., M.; Weston, S.; Balhorn, R. (1981). DNA and protein content of mouse sperm. Implications regarding sperm chromatin structure. Exp Cell Res 136

Punjani, A., Rubinstein, J. L., Fleet, D. J, Brubaker, M. A. (2017). cryoSPARC: algorithms for rapid unsupervised cryo-EM structure determination. Nat Methods 14

Qian, M. X., Pang, Y., Liu, C. H.,…Qiu, X.B. (2013). Acetylation-mediated proteasomal degradation of core histones during DNA repair and spermatogenesis. Cell 153, 1012–24.

Ram, A., Jalal, S., Jalal, A. S. and Kumar, M. (2010). A Density based Algorithm for Discovering Density Varied Clusters in Large Spatial Databases. International Journal of Computer Applications 3

Rathke, C., Baarends, W. M.; Awe, S.; Renkawitz-Pohl, R. (2014). Chromatin dynamics during spermiogenesis. Biochim Biophys Acta 1839

Rawe, V. Y., Diaz, E. S., Abdelmassih, R.,…Chemes, H.E. (2008). The role of sperm proteasomes during sperm aster formation and early zygote development: implications for fertilization failure in humans. Hum Reprod 23, 573–80.

Rohou, A., Grigorieff, N. (2015). CTFFIND4: Fast and accurate defocus estimation from electron micrographs. J Struct Biol 192

Russo, C. J., Scotcher, S. and Kyte, M. (2016). A precision cryostat design for manual and semi-automated cryo-plunge instruments. Rev Sci Instrum 87, 114302.

Saldivar-Hernandez, A., Gonzalez-Gonzalez, M. E., Sanchez-Tusie, A.,…Chirinos, M. (2015). Human sperm degradation of zona pellucida proteins contributes to fertilization. Reprod Biol Endocrinol 13, 99.

Santos, Á. d., Knowles, O., Dendooven, T.,…Allegretti, M. (2024). Human spermatogenesis leads to a reduced nuclear pore structure and function. BioRxiv

Sato, B., Kim, J., Morohoshi, K.,…Chiba, T. (2023). Proteasome-Associated Proteins, PA200 and ECPAS, Are Essential for Murine Spermatogenesis. Biomolecules 13

Schertel, A., Snaidero, N., Han, H. M.,…Mobius, W. (2013). Cryo FIB-SEM: volume imaging of cellular ultrastructure in native frozen specimens. J Struct Biol 184, 355–60.

Schindelin, J., Arganda-Carreras, I., Frise, E.,…Cardona, A. (2012). Fiji: an open-source platform for biological-image analysis. Nat Methods 9, 676–82.

Schrader J., H. F., Mata R. A., Tittmann K., Schneider T. R., Stark H., Bourenkov G., Chari A. (2016). The inhibition mechanism of human 20S proteasomes enables next-generation inhibitor design. Science 5

Schweitzer, A., Aufderheide, A., Rudack, T.,…Baumeister, W. (2016). Structure of the human 26S proteasome at a resolution of 3.9 A. Proc Natl Acad Sci U S A 113, 7816–21.

Smith, D. M., Chang, S. C., Park, S., Finley, D., Cheng, Y., Goldberg, A. L. (2007). Docking of the proteasomal ATPases’ carboxyl termini in the 20S proteasome’s alpha ring opens the gate for substrate entry. Mol Cell

Song, W. H., Zuidema, D., Yi, Y. J.,…Sutovsky, P. (2021). Mammalian Cell-Free System Recapitulates the Early Events of Post-Fertilization Sperm Mitophagy. Cells 10

Sutovsky, P. (2011). Sperm proteasome and fertilization. Reproduction 142, 1–14.

Sutovsky, P., Manandhar, G., McCauley, T. C.,…Day, B.N. (2004). Proteasomal interference prevents zona pellucida penetration and fertilization in mammals. Biol Reprod 71, 1625–37.

Tanaka, A., Nagayoshi, M., Tanaka, I. and Kusunoki, H. (2012). Human sperm head vacuoles are physiological structures formed during the sperm development and maturation process. Fertil Steril 98, 315–20.

Tegunov, D. and Cramer, P. (2019). Real-time cryo-electron microscopy data preprocessing with Warp. Nat Methods 16, 1146–1152.

Tegunov, D., Xue, L., Dienemann, C., Cramer, P. and Mahamid, J. (2021). Multi-particle cryo-EM refinement with M visualizes ribosome-antibiotic complex at 3.5 A in cells. Nat Methods 18, 186–193.

Toste Rego, A. and da Fonseca, P. C. A. (2019). Characterization of Fully Recombinant Human 20S and 20S-PA200 Proteasome Complexes. Mol Cell 76, 138–147 e5.

Uriarte, M., Sen Nkwe, N., Tremblay, R.,…Affar, E.B. (2021). Starvation-induced proteasome assemblies in the nucleus link amino acid supply to apoptosis. Nat Commun 12, 6984.

Ustrell, V., Hoffman, L., Pratt, G., Rechsteiner, M. (2002). PA200, a nuclear proteasome activator involved in DNA repair. EMBO J 21

Voges, D. Z. P.; Baumeister, W. (1999). The 26S proteasome: a molecular machine designed for controlled proteolysis. Annu Rev Biochem 68

Wagner, T., Merino, F., Stabrin, M.,…Raunser, S. (2019). SPHIRE-crYOLO is a fast and accurate fully automated particle picker for cryo-EM. Commun Biol 2, 218.

Wang, L., Liu, C., Wang, X.,…Li, W. (2025). Separately prestored proteasome components to prevent polyspermy. BioRviv

Williams, C. J., Headd, J. J., Moriarty, N. W.,…Richardson, D.C. (2018). MolProbity: More and better reference data for improved all-atom structure validation. Protein Sci 27, 293–315.

Wolf, M., DeRosier, D. J., Grigorieff, N. (2006). Ewald sphere correction for single-particle electron microscopy. Ultramicroscopy 106

Xin, B. T., Huber, E. M., de Bruin, G.,…Overkleeft, H.S. (2019). Structure-Based Design of Inhibitors Selective for Human Proteasome beta2c or beta2i Subunits. J Med Chem 62, 1626–1642.

Xiong, Y., Yu, C. and Zhang, Q. (2022). Ubiquitin-Proteasome System-Regulated Protein Degradation in Spermatogenesis. Cells 11

Yasuda, S., Tsuchiya, H., Kaiho, A.,…Saeki, Y. (2020). Stress- and ubiquitylation-dependent phase separation of the proteasome. Nature 578, 296–300.

Zaffagnini, G., Cheng, S., Salzer, M. C.,…Boke, E. (2024). Mouse oocytes sequester aggregated proteins in degradative super-organelles. Cell 187, 1109–1126 e21.

Zhang, H., Zhou, C., Mohammad, Z. and Zhao, J. (2024). Structural basis of human 20S proteasome biogenesis. Nat Commun 15, 8184.

Zhang, M., Chiozzi, R. Z., Skerrett-Byrne, D. A.,…Bromfield, E. G. (2022). High Resolution Proteomic Analysis of Subcellular Fractionated Boar Spermatozoa Provides Comprehensive Insights Into Perinuclear Theca-Residing Proteins. Front Cell Dev Biol 10, 836208.

Zhang, Q., Ji, S. Y., Busayavalasa, K., Shao, J. and Yu, C. (2019). Meiosis I progression in spermatogenesis requires a type of testis-specific 20S core proteasome. Nat Commun 10, 3387.

Zhang, X., Yang, J., Yang, W., Cui, N., Duan, T., Li, S., Cao, J., Bush, S. J., Tong, G. (2025). Discordant effects of maternal age on the human MII oocyte transcriptome. Mol Hum Reprod

Zhang, Z. H., Jiang, T. X., Chen, L. B.,…Qiu, X.B. (2021). Proteasome subunit alpha4s is essential for formation of spermatoproteasomes and histone degradation during meiotic DNA repair in spermatocytes. J Biol Chem 296, 100130.

Zheng, S., Wolff, G., Greenan, G.,…Agard, D.A. (2022). AreTomo: An integrated software package for automated marker-free, motion-corrected cryo-electron tomographic alignment and reconstruction. J Struct Biol X 6, 100068.

Zheng, S. Q., Palovcak, E., Armache, J. P.,…Agard, D.A. (2017). MotionCor2: anisotropic correction of beam-induced motion for improved cryo-electron microscopy. Nat Methods 14, 331–332.

Zigo, M., Manaskova-Postlerova, P., Jonakova, V., Kerns, K. and Sutovsky, P. (2019). Compartmentalization of the proteasome-interacting proteins during sperm capacitation. Sci Rep 9, 12583.

Zimmerman, S. and Sutovsky, P. (2009). The sperm proteasome during sperm capacitation and fertilization. J Reprod Immunol 83, 19–25.

Zimmerman, S. W., Manandhar, G., Yi, Y. J.,…Sutovsky, P. (2011). Sperm proteasomes degrade sperm receptor on the egg zona pellucida during mammalian fertilization. PLoS One 6, e17256.

Zivanov, J., Nakane, T. and Scheres, S. H. W. (2019). A Bayesian approach to beam-induced motion correction in cryo-EM single-particle analysis. IUCrJ 6, 5–17.

Zivanov, J., Nakane, T. and Scheres, S. H. W. (2020). Estimation of high-order aberrations and anisotropic magnification from cryo-EM data sets in RELION-3.1. IUCrJ 7, 253–267.

Zivanov, J., Oton, J., Ke, Z.,…Scheres, S. H.W. (2022). A Bayesian approach to single-particle electron cryo-tomography in RELION-4.0. Elife 11

Zivkovic, D., Sanchez Dafun, A., Menneteau, T.,…Bousquet, M.P. (2022). Proteasome complexes experience profound structural and functional rearrangements throughout mammalian spermatogenesis. Proc Natl Acad Sci U S A 119, e2116826119.

Zuiderveld, K. (1994). Contrast limited adaptive histogram equalization. Graphics gems IV

